# Interspecies blastocyst complementation generates functional rat cell-derived forebrain tissues in mice

**DOI:** 10.1101/2023.04.13.536774

**Authors:** Jia Huang, Bingbing He, Xiali Yang, Xin Long, Yinghui Wei, Yanxia Gao, Yuan Fang, Wenqin Ying, Zikang Wang, Chao Li, Yingsi Zhou, Shuaishuai Li, Linyu Shi, Fan Guo, Haibo Zhou, Hui Yang, Jun Wu

**Author notes:** Correspondence (F.G.), (H.Z.), (H.Y.), (J.W.). These authors contributed equally to this work (Jia Huang, Bingbing He, Xiali Yang, Xin Long, Yinghui Wei).

## Abstract

Interspecies organogenesis via blastocyst complementation provides a unique platform to study development in an evolutionarily context and holds potential to overcome world-wide organ shortages^1^. By using this technique, rat pancreas, thymus, heart, and eye tissues have been generated in mice^2–4^. To date, however, xeno-generation of brain tissues has not been achieved through blastocyst complementation. Here, we developed an optimized one-step blastocyst complementation strategy based on C-CRISPR^5^, which facilitated rapid screening of candidate genes to support blastocyst complementation. Among the seven WNT pathway-related genes selected for targeting, only *Dkk1* or *Hesx1* deficiency supported forebrain complementation by blastocyst injection of mouse embryonic stem cells (mESCs). Further, injecting rat ESCs (rESCs) into mouse blastocysts deficient for *Hesx1* but not *Dkk1* supported the development of adult chimeric forebrains comprised a large proportion of rat cells that were structurally and functionally similar to the mouse forebrains. Our analysis revealed that the rESC-derived forebrains developed along the spatial-temporal trajectory with the mouse forebrains rather than rat forebrains, but gene expression profiles of rESC-derived nerve cells surprisingly maintained the characteristics of the rat cells. We noted that the chimeric rate gradually decreased as development progressed, suggesting xenogeneic barriers during mid-to-late prenatal development. Interspecies forebrain complementation opens the door for studying evolutionarily conserved and divergent mechanisms underlying brain development and cognitive function. The C-CRIPSR based IBC strategy developed here holds great potential to broaden the study and application of interspecies organogenesis.

## INTRODUCTION

Advances in interspecies chimeras and blastocyst complementation have offered hope in addressing the global shortage of donor organs and have expanded our understanding of the molecular and cellular mechanisms involved in organogenesis. Blastocyst complementation involves injecting chimera-competent donor pluripotent stem cells (PSCs) into mutant host blastocysts, which lack one or more essential genes for the development of a specific organ. Through intra- or inter-species chimera formation, donor PSCs can fill the vacant developmental organ niche and generate organs derived from donor PSCs within the host^6, 7^. Previous studies have successfully demonstrated intra-species blastocyst complementation for various mouse tissues, including the pancreas, thymus, kidney, heart, liver, lung, and forebrain^8–19^. Although limited in scope, several studies have attempted inter-species blastocyst complementation, generating rat pancreas, thymus, hematoendothelial tissues, and germ cells in mice^2–4, 20–22^, as well as mouse pancreas, kidney, and germ cells in rats^23–25^. To date, however, inter-species blastocyst complementation has not been achieved for brain tissues. There is substantial interest in generating brain tissues from one species within another, as this would not only enable in vivo studies of brain development and function in an evolutionary context, but also provide a crucial foundation for addressing ethical concerns surrounding the contribution of human PSCs to animal brains1^1^.

Traditional blastocyst complementation methods involve producing sexually mature gene-edited mice^4, 11^, and breeding these mice can be time-consuming and labor-intensive. The majority of newborn pups are not complemented, necessitating the inspection of each pup individually to identify successful blastocyst replacement. This process can take a considerable amount of time (10-26 months) to determine whether a selected candidate gene is suitable for the target organ’s blastocyst complementation. Unlike several other tissues, suitable gene(s) for interspecies neural blastocyst complementation have yet to be identified. Screening candidate genes through targeted gene disruption in germline competent PSCs is a time-consuming process, even for species with short gestation periods and rapid sexual maturity. This approach becomes impractical for large livestock species and non-human primates. In this study, we present an optimized blastocyst complementation method that enables rapid screening of candidate genes and facilitates the generation of functional rat embryonic stem cell (rESC)-derived forebrain tissues in mice.

## RESULTS

### CCBC, an improved blastocyst complementation method

To overcome the limitations associated with existing blastocyst complementation methods, we combined the <u>C</u>-<u>C</u>RISPR approach, which enables rapid screening of candidate genes and one-step generation of complete gene knockout animals with multiple sgRNAs^5^, with <u>b</u>lastocyst <u>c</u>omplementation (CCBC) (Figure 1A left). CCBC allows for quick testing whether the target genes are suitable for blastocyst complementation and enables one-step generation of organ-reconstituted chimeras, which is desirable when using large livestock hosts.

**Figure 1.**
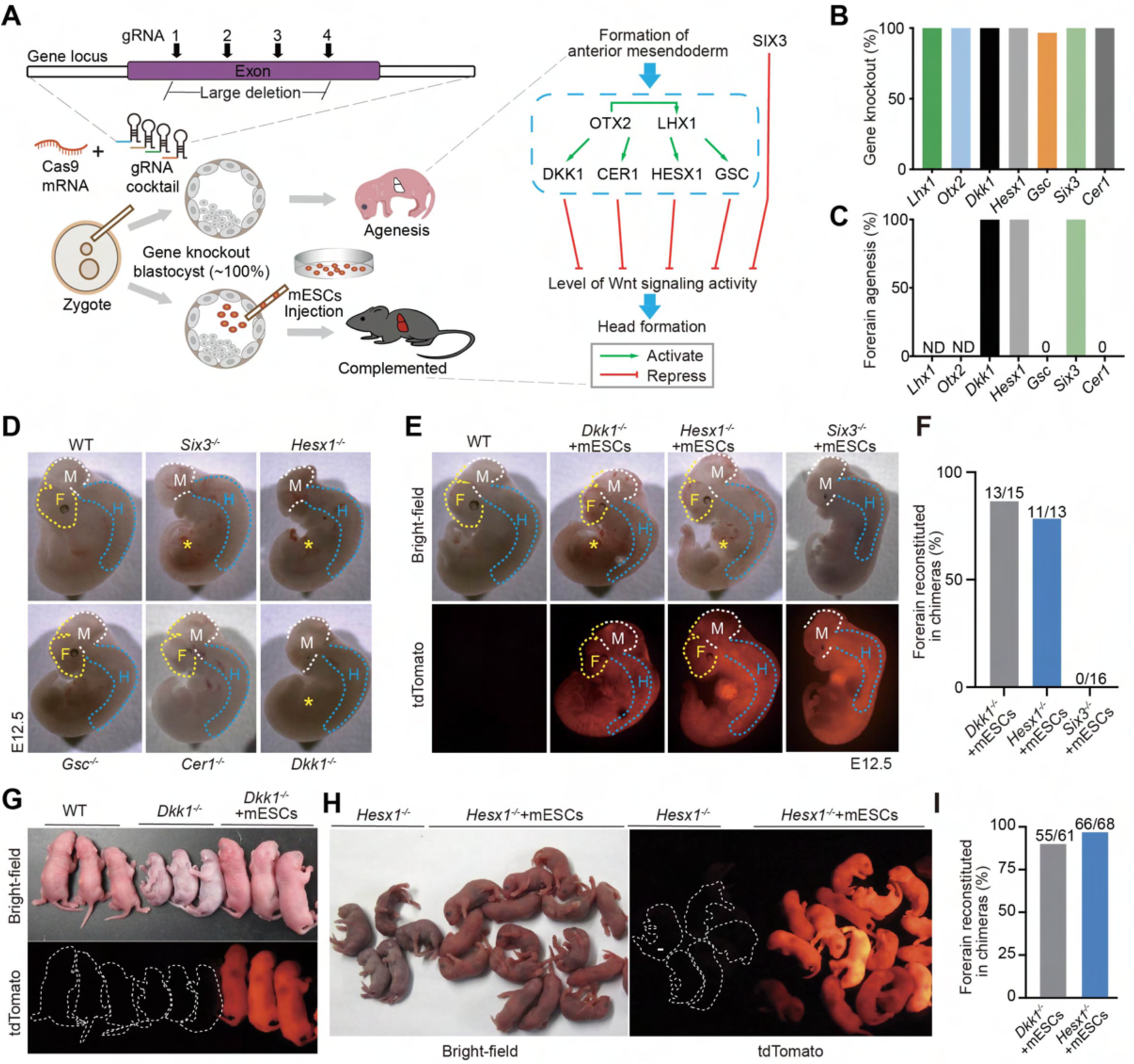
CCBC facilitates quick genetic screening for neural blastocyst complementation. (A) Schematic showing one-step generation specific organ in mice using CCBC and candidate genes for forebrain compensation which involved in regulating Wnt signaling during brain organogenesis. (B) The efficiency of C-CRISPR-mediated gene knockout in 4-cell mouse embryos. (C) The percentage of forebrain agenesis after gene knockout in E12.5 mouse embryos. ND, no data. (D) Representative images showing forebrain agenesis at embryonic day (E) 12.5 in gene knockout mouse embryos. F, forebrain; M, midbrain; H, hindbrain. Experiments were repeated 3 times independently for each group. (E) Representative images showing forebrain reconstitution in E12.5 CCBC mouse embryos. F, forebrain; M, midbrain; H, hindbrain. (F) The efficiency of forebrain reconstitution in E12.5 CCBC chimeras. (G) Images showing newborn *Dkk1^-/-^* mice (P0) with or without forebrain reconstitution. Note that the complete allogenic forebrain was only formed in *Dkk1^-/-^*+mESCs chimeras but not in *Dkk1^-/-^*mice. (H) Images showing newborn *Hesx1^-/-^* mice (P0) with or without brain reconstitution. Note that the forebrain was only formed in *Hesx1^-/-^*+mESCs chimera but not in *Hesx1^-/-^* mice. (I) Quantification of forebrain reconstitution efficiency of *Dkk1^-/-^*+mESCs and *Hesx1^-/-^*+mESCs chimeras at P0.

The transcription factor *Pdx1* plays a critical role in pancreatic development: mice lacking *Pdx1* do not develop a pancreas and die soon after birth^26^. Previous studies have shown that disruption of *Pdx1* gene in the host embryos can support mouse-mouse, rat-mouse and mouse-rat pancreas blastocyst complementation^2, 4, 15, 25^. Using pancreas as a proof-of-concept, we tested the efficacy and efficiency of CCBC (Figure S1A;). We designed 4 sgRNAs that are spaced out 30-55bp apart from each other within the exon 1 of the mouse *Pdx1* locus (Figure S1B) and co-injected them with Cas9 mRNA into mouse zygotes. We found *Pdx1* was successfully disrupted from all the examined 4-cell embryos using the C-CRISPR approach (Figure S1C). All neonatal *Pdx1*^-/-^ mice generated using CCBC died within 4 days after birth due to pancreas agenesis (Figures S1D-S1F).

For pancreas blastocyst complementation, 4 gRNAs (100ng/μl) targeting *Pdx1* and Cas9 mRNA (80ng/μl) were co-injected into mouse zygotes followed by injection of 8-10 tdTomato labeled mESCs into blastocysts. Among the 37 *Pdx1*^-/-^ mice that were born, 14 were tdTomato positive among which 13 showed an intact pancreas; the remaining pups displayed pancreatic agenesis (Figures S1G-S1I; Table S1). In contrast to the WT+mESCs chimeras, we found that the pancreas of *Pdx1^-/-^*+mESCs were mostly derived from donor mESCs (Figure S1J), consistent with a previous study.^4^ Immunofluorescence analysis demonstrated that the pancreas of *Pdx1^-/-^*+mESCs chimeras contained cells expressing the pancreatic exocrine protein α-AMYLASE and the pancreatic endocrine proteins INSULIN and GLUCAGON, which were completely overlapped with tdTomato signals (Figure S1K).

### CCBC facilitates quick genetic screening for neural blastocyst complementation

Leveraging the Cre-DTA (diphtheria toxin A) mediated cell ablation, a recent study harnessed blastocyst complementation for allogenic forebrain organogenesis in mice^11^. To date, it remains unknown which gene(s) is suitable to target for use in gene knockout-based forebrain blastocyst complementation. In early vertebrate embryogenesis, head formation is known to depend on the Wnt/β-catenin signaling^27–31^. To this end, we used C-CRISPR to perform a targeted screen of seven known genes that are implicated in regulating the canonical Wnt signaling during brain development (Figure 1A right).

All seven selected candidate genes could be efficiently deleted by C-CRISPR (Figure 1B), but only knockout of *Dkk1*, *Hesx1*, and *Six3* resulted in a forebrain agenesis phenotype (Figures 1C and 1D). Upon injection of mESCs into individual gene-deleted blastocysts, we observed complete allogenic forebrains in *Dkk1^-/-^* and *Hesx1^-/-^* E12.5 embryos (Figures 1E and 1F). In contrast, mESCs were not able to rescue the forebrain agenesis phenotype in *Six3^-/-^* E12.5 embryos (Figures 1E and 1F). Among the 61 *Dkk1*^-/-^+mESCs newborn chimeras, we obtained 55 (90.16%) forebrain reconstituted chimeras (Figures 1G and 1I). Among the 68 *Hesx1*^-/-^+mESCs newborn chimeras, we obtained 66 (50.77%) forebrain reconstituted chimeras (Figures 1H and 1I). Whole genome sequencing of *Hesx1*^-/-^ mice (n = 6) showed no off-target effects were detected (Table S2).

To determine the contribution of donor cells in the chimeric forebrains, we injected tdTomato^+^ mESCs into *Hesx1*^-/-^ blastocysts expressing EGFP driven by the ubiquitous promoter CAG (Figure 2A), and generated EGFP-tdTomato dual-labelled chimeras (Figure 2B). Analysis of whole brain sagittal slices revealed most, if not all, cells in the cerebral cortex and hippocampus of the *Hesx1*^-/-^+mESCs chimeras were tdTomato^+^, and the proportion of tdTomato^+^ within forebrain was much higher in *Hesx1*^-/-^+mESCs chimeras (∼ 100%) than in WT+mESCs chimeras (∼50%) (Figures 2C-E; Figure S3A). The midbrain area unaffected by *Hesx1* knockout had a chimeric rate close to 50% (Figure 2E; Figures S2A and S2B). These findings show complete repopulation of the vacant host forebrain niche by donor mESCs. By contrast, *Dkk1*^-/-^+mESCs chimeras showed on average 95.93% and 98.76% chimeric rates in the midbrain and cortex while 53.36% in the hippocampus (Figures S2A and S2B). Thus, it appears that *Dkk1* is not suitable for forebrain blastocyst complementation. All *Hesx1*^-/-^+mESCs and *Dkk1*^-/-^+mESCs mice survived to adulthood (Figure 2F and Figure S2C), with brain size comparable to WT-mESCs controls (Figures 2G and 2H; Figures S2D and S2E). Taken together, these results demonstrate the efficacy of using CCBC to screen and identify candidate genes for blastocyst complementation and show the proof-of-concept of targeting *Hesx1* for intraspecies functional forebrain complementation in mice.

**Figure 2.**
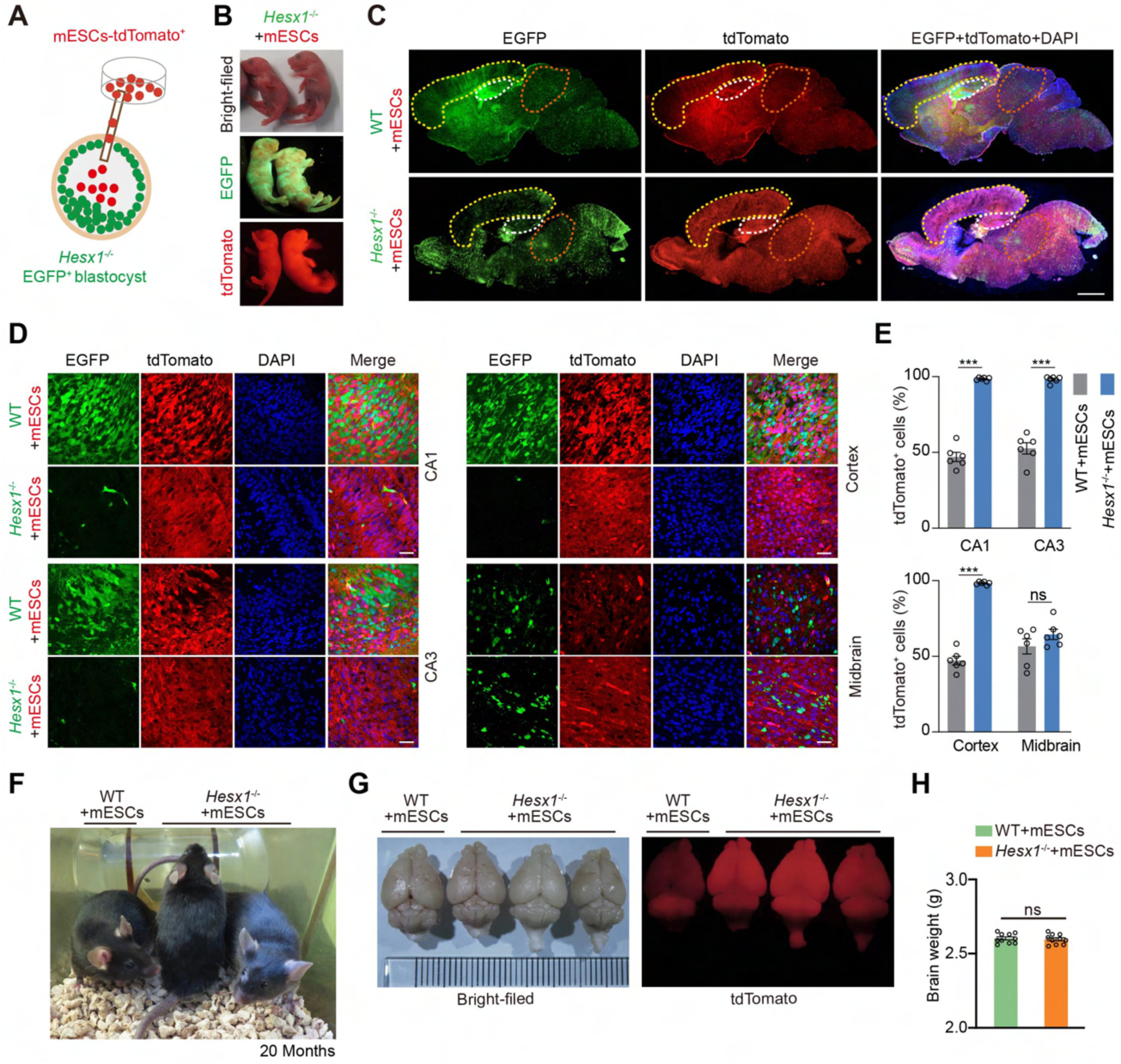
Intraspecies blastocyst complementation of forebrains in *Hesx1*^-/-^ mice. (A) Schematic for the generation of dual-color-labelled chimeric mice. Red, tdTomato^+^ donor mESCs; Green, EGFP^+^ *Hesx1^-/-^*host blastocyst. (B) Representative images of *Hesx1*^-/-^+mESCs and WT+mESCs chimeras at P0. (C) Representative sagittal brain sections from WT+mESCs (lower, n = 3) and *Hesx1*^-/-^+mESCs (upper, n = 4) chimeras. Yellow dashed line, cortex; white dashed line, hippocampus; orange dashed line, midbrain. Scale bars, 1mm. (D) Representative images showing the contribution of tdTomato^+^ donor mESCs to the CA1, CA3, cortex and midbrain regions in WT+mESCs (upper, n = 3) and *Hesx1*^-/-^+mESCs (lower, n = 4) P0 chimeras. Green, host-derived cells; Red, donor-derived cells; Blue, DAPI-stained nuclei. Scale bars, 100 μm. (E) Percentages of tdTomato^+^ donor mouse cells in the CA1, CA3, cortex and midbrain regions in WT+mESCs and *Hesx1*^-/-^+mESCs chimeras (n = 3 per group). All values are presented as the mean ± s.e.m.. ***p < 0.001, unpaired t-tests. ns, not significant. (F) *Hesx1^-/-^*+mESCs and WT+mESCs chimeras at the age of 20 months. (G) Bright-field and fluorescence images showing no obvious difference between the brains of *Hesx1^-/-^*+mESCs and WT+mESCs chimeras at 10 weeks of age. (H) Brain weight of *Hesx1^-/-^*+mESCs and WT+mESCs at 10 weeks of age (n = 10 mice per group).

### CCBC enables interspecies neural blastocyst complementation

To determine whether CCBC is applicable to interspecies blastocyst complementation, we first injected 4 sgRNAs (100ng/μl) targeting *Pdx1* along with Cas9 mRNA (80ng/μl) into mouse zygotes followed by blastocyst injection of 8-10 tdTomato^+^ rESCs. Among the 50 rat-mouse chimeras born, we obtained 9 bearing a reconstituted rat pancreas (Figures S3A and S3B). Pancreatic cells in reconstituted *Pdx1*^-/-^+rESCs chimeras expressed α-AMYLASE, INSULIN, and GLUCAGON, and were mostly derived from rESCs (Figure S3C), which is consistent with previous reports^2, 4^. These results show that a xenogeneic rat pancreas can be successfully formed in *Pdx1*^-/-^ mice by combining C-CRISPR and interspecies blastocyst complementation, demonstrating the utility of the CCBC method for one-step generation of xenogeneic organs and tissues.

To date, interspecies blastocyst complementation has not been reported for any brain tissues. To determine whether rat forebrain tissues can be generated in mice via CCBC, we injected rESCs into *Dkk1*^-/-^*or Hesx1*^-/-^ mouse blastocysts (Figure 3A). In total, 417 *Hesx1*^-/-^+rESCs chimeras were born, among which 401 were forebrain partial reconstituted (96.16%) and 16 were forebrain reconstituted (3.84%) (Figures 3B and 3C). In contrast, all born *Dkk1*^-/-^+rESCs chimera pups (374) were forebrain partial reconstituted (100%) and no forebrain reconstituted (0%) (Figures S4A and S4B). We found that, in the *Hesx1*^-/-^+rESCs chimeras, the proportion of tdTomato^+^ cells were higher in the cortex and hippocampus than in other brain regions and other tissues (Figures 3D-3G; Figures S4C and S4D).

**Figure 3.**
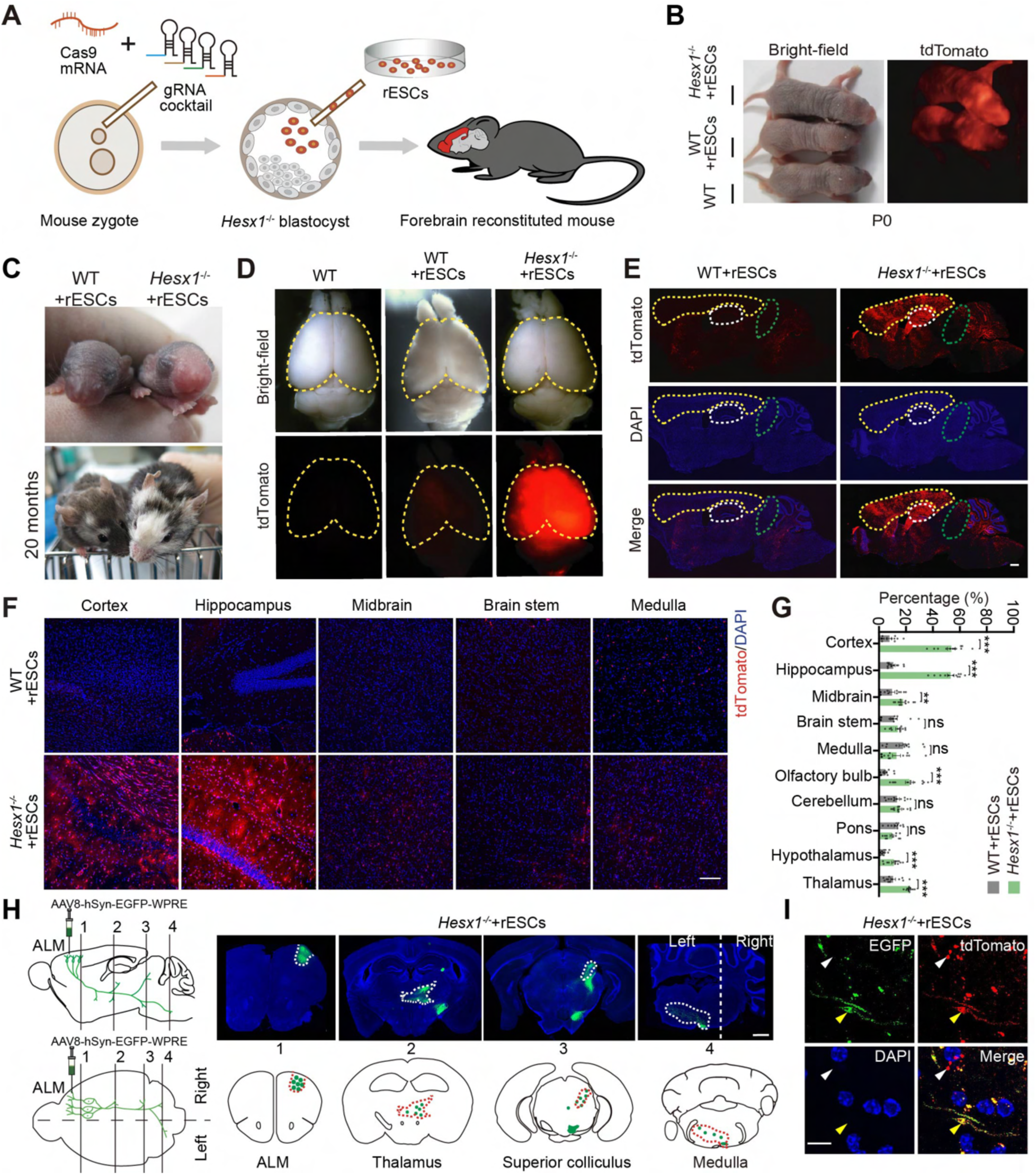
Generation of rat forebrain tissues in mice via interspecies blastocyst complementation. (A) Schematic for the generation of rESC-derived forebrain tissues in mice via CCBC. (B) Bright-field and fluorescence images of WT mice, WT+mESCs and *Hesx1*^-/-^+rESCs chimeras at P0. (C) Images of WT+rESCs and *Hesx1*^-/-^+rESCs chimeras at P3 (top) and 20 months (bottom). (D) Images of brains from a WT mouse, WT+rESCs, and *Hesx1*^-/-^+rESCs chimeras at 10 weeks. (E) Sagittal brain sections from WT+rESCs (left) and *Hesx1*^-/-^+rESCs (right) chimeras at 10 weeks. Yellow dashed line, cortex; white dashed line, hippocampus; green dashed line, midbrain. Scale bar, 1mm. (F) Representative images showing the contribution of tdTomato^+^ rat cells to different brain regions in WT+rESCs and *Hesx1^-/-^*+rESCs chimeras (n = 3 per group). (G) Percentages of tdTomato^+^ rat cells in the indicated brain regions in WT+rESCs and *Hesx1*^-/-^ +rESCs chimeras (n = 3 per group). Unpaired t-tests. (H) The axonal projections of neurons from a forebrain region anterior lateral motor cortex (ALM). Note that strong signals were observed in the cortex, thalamus, superior colliculus (SC), and medulla, the known target regions of ALM neuron axons (n = 2 chimeras). Scale bar, 1 mm. (I) Images from a coronal slice showing that the axon fibers of ALM neurons in the midbrain were positive for both EGFP and tdTomato, suggesting that rESC-derived tdTomato^+^ neurons send axons to the midbrain. Yellow arrowheads indicate the colocalization of tdTomato and EGFP signals in the axonal fiber. White arrowheads indicate projected host-derived axonal fibers. Scale bar, 10 μm. All values are presented as the mean ± s.e.m.. **p < 0.01, ***p < 0.001, unpaired t-tests. ns, not significant.

To test whether the neurons derived from rESCs can correctly project and make neuronal connections in rat-mouse chimeric brains, we injected AAV8-hSyn-EGFP into the anterior lateral motor cortex (ALM) of the *Hesx1*^-/-^+rESCs forebrain. Consistent with a previous report,^32^ we found that EGFP^+^ axons sent axon projections from ALM to thalamus, superior colliculus and medulla areas in the midbrain and brain stem (Figures 3H and 3I), suggesting that rESC-derived forebrain neurons formed the expected connections with other brain regions. Taken together, these results demonstrate that *Hesx1*^-/-^ mouse blastocysts provide a developmental niche amenable to the formation of functional rat forebrain tissues from donor rESCs.

### Forebrains reconstituted in intra- or interspecies chimeras are structurally and functionally normal

All *Hesx1*^-/-^+rESCs chimeras survived to adulthood showed a body weight curve comparable to that of WT+mESCs and *Hesx1*^-/-^+mESCs chimeras (Figure 4A). The structural integrity of rat or mouse forebrain tissues generated in mice was assessed by layer-specific immunocytochemical marker staining. The cortex layer V and hippocampus of WT+mESCs, *Hesx1*^-/-^+mESCs and *Hesx1*^-/-^+rESCs chimeras contained cells expressing the neural marker Ctip2 (Figure 4B). And both the thickness and cell density of various layers of the cerebral cortex and the hippocampus appeared similar among *Hesx1*^-/-^+rESCs, WT+mESCs and *Hesx1*^-/-^+mESCs chimeras (Figures 4C-4E). These results are consistent with previous observations that body size and xenogeneic organ size matched with the host species in interspecies chimeras^2, 4, 25^.

**Figure 4.**
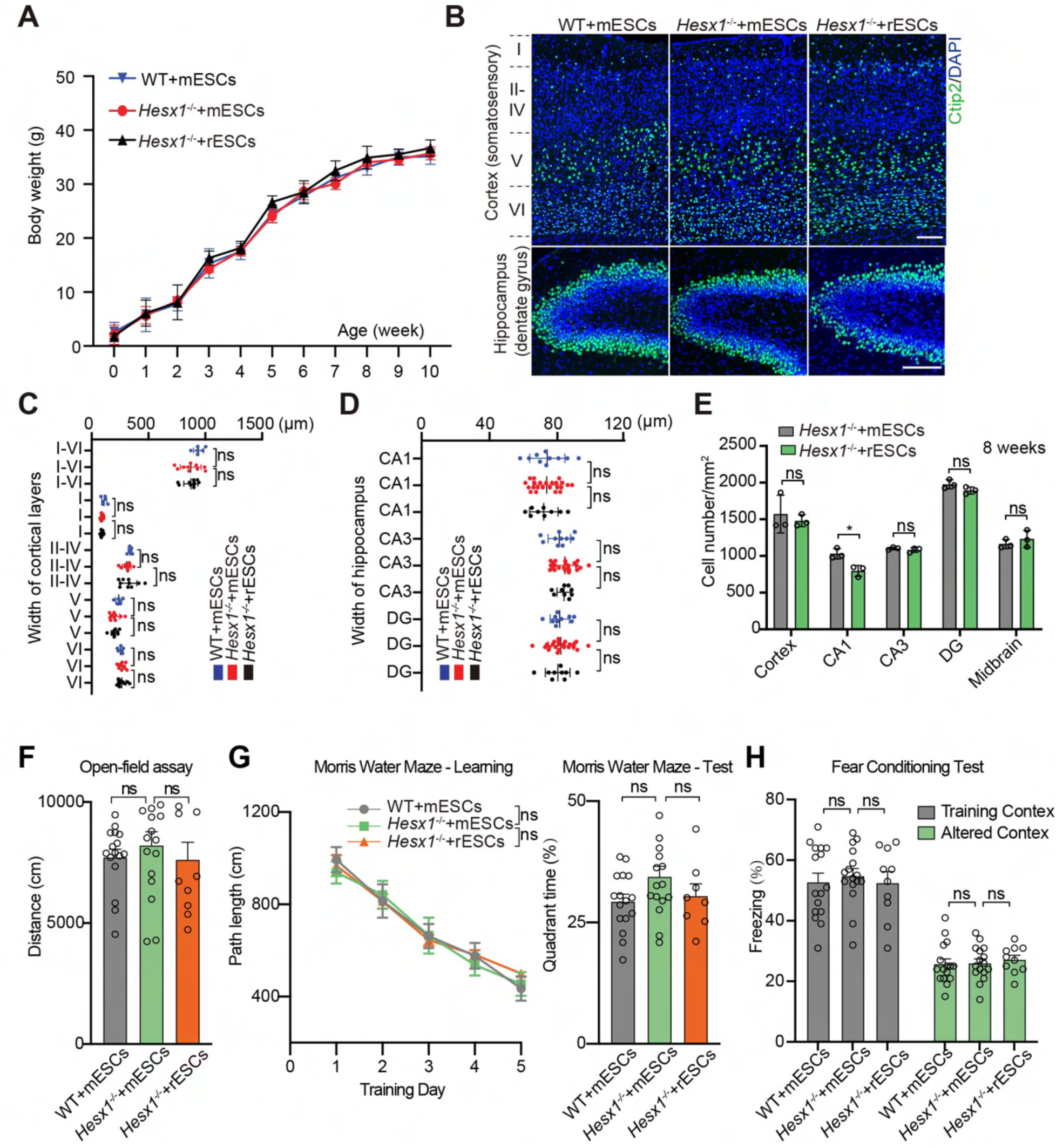
Reconstituted forebrains of intra- or interspecies chimeras are structurally and functionally normal. (A) Body weight curves of *Hesx1^-/-^*+rESCs, *Hesx1^-/-^*+mESCs and WT+mESCs chimeras. (n = 10 per group). (B) Representative images of sections from the somatosensory cortex and the hippocampus stained with Ctip2 and DAPI from WT+mESCs (left) and *Hesx1*^-/-^+mESCs (right) chimeras at P7. Scale bar, 100 μm. (C) Widths of somatosensory cortex layers in WT+mESCs (blue, n = 3), *Hesx1*^-/-^+mESCs (red, n = 5), or *Hesx1*^-/-^+rESCs (black, n = 5) chimeras at P7. (D) Widths of the dentate gyrus (DG), CA3, and CA1 regions in WT+mESCs (blue, n = 3), *Hesx1*^-/-^+mESCs (red, n = 5), or *Hesx1*^-/-^+rESCs (black, n = 5) chimeras at P7. (E) Quantitative analysis of cell density of different brain regions of *Hesx1*^-/-^+mESCs and *Hesx1*^-/-^+rESCs chimeras at the age of 8 weeks (n = 3 per group). (F) Total distance travelled in the open-field assay for WT+rESCs (gray), *Hesx1*^-/-^+mESCs (green), and *Hesx1*^-/-^+rESCs (orange) chimeras. (G) Left. Mean path length to the platform for WT+rESCs (gray), *Hesx1*^-/-^+mESCs (green), and *Hesx1*^-/-^+rESCs (orange) chimeras in the learning trials of Morris water maze task. Right. Mean time spent in the target quadrant in the target quadrant for WT+rESCs (gray), *Hesx1*^-/-^+mESCs (green), and *Hesx1*^-/-^+rESCs (orange) chimeras in test trials of Morris water maze task. (H) Percentage of freezing of WT+rESCs, *Hesx1*^-/-^+mESCs, and *Hesx1*^-/-^+rESCs chimeras in the contextual fear conditioning test. For Fig. 4F-4H, n = 16 chimeras for the WT+mESCs and *Hesx1^-/-^* +mESCs groups; n = 10 chimeras for the *Hesx1^-/-^*+rESCs group. All values are presented as the mean ± s.e.m.. **p < 0.01, ***p < 0.001, unpaired t-tests. ns, not significant.

We also assessed cognitive functions using behavioral tests known to depend on intact forebrain function, which include the Morris water maze, Open-field assay, and Contextual fear conditioning^33–35^. We observed no differences in performance between WT+mESCs, *Hesx1*^-/-^ +rESCs and *Hesx1*^-/-^+mESCs chimeras (Figures 4F-4H) as well as between WT+mESCs and *Dkk1*^-/-^+mESCs chimeras (Figures S2F-S2I), for any of these tests, suggesting normal functions of the reconstituted forebrains.

### Xenogeneic barriers during mid-to-late stage of embryonic development

Whole embryos sections of embryos at different stages of development were performed to monitor the dynamic donor rat cell contribution in *Hesx1^-/-^*+rESCs embryos and fetsues. In agree with a previous report^36^, we observed that as the development progressed, the overall chimeric contribution of rat cells in mouse fetuses gradually decreased from ∼60% to about 20% by E17.5, and the rat chimeric level in the forebrain also decreased from 90-100% to ∼60% by E15.5 (Figures 5A-5C). These results suggest an interspecies barrier from mid-to-late gestation stages between rats and mice, and to enable full forebrain complementation of *Hesx1^-/-^* mouse by rat cells, method(s) to overcome chimeric rate reduction during this developmental period need to be considered^37, 38^.

**Figure 5.**
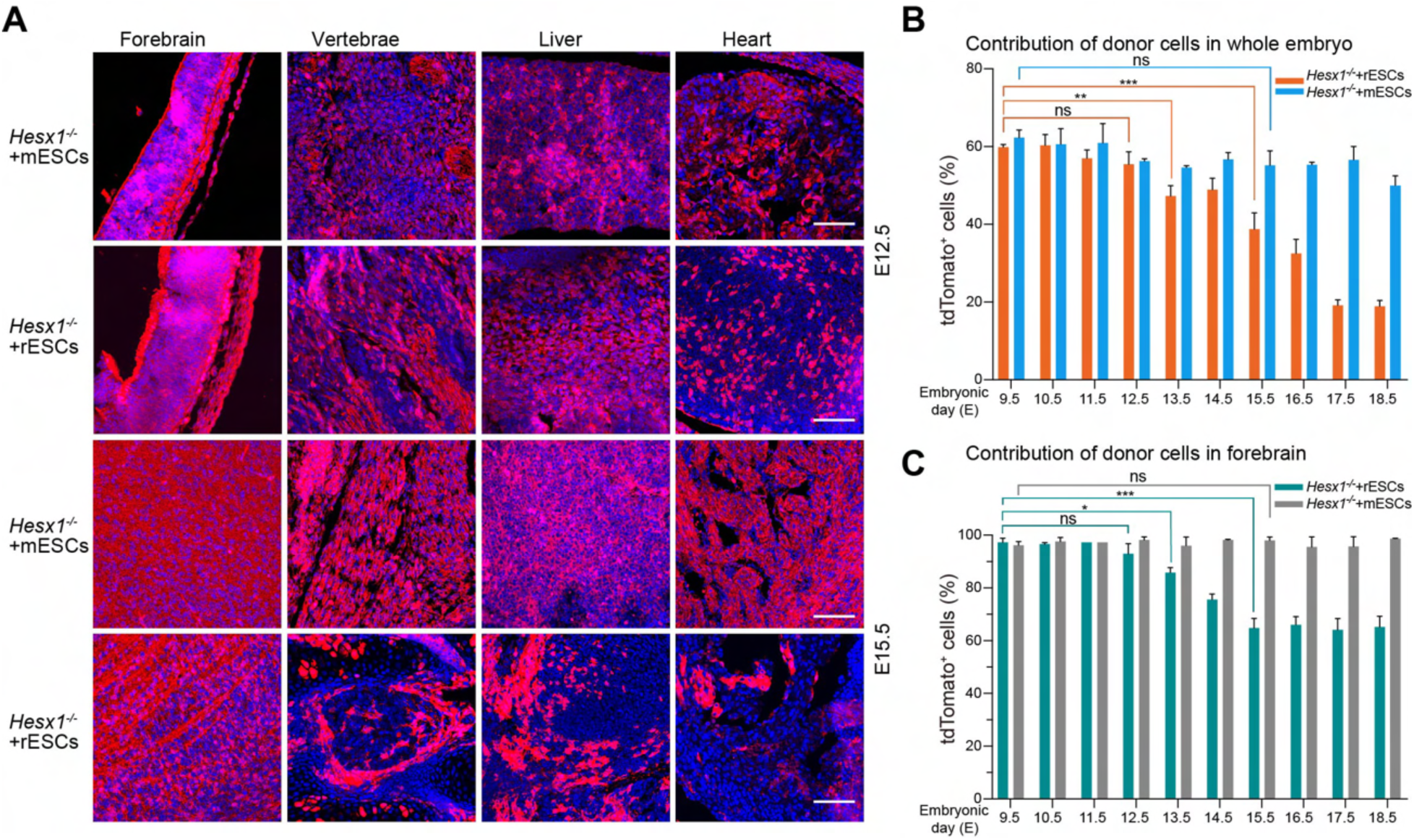
The dynamic donor rat cell contribution in *Hesx1^-/-^*+rESCs embryos and fetsues. (A) Representative fluorescent images of different tissue sections in *Hesx1^-/-^*+mESCs and *Hesx1^-/-^*+rESCs fetuses at E12.5 and E15.5. Scale bars, 100 μm. (B) and (C) Contribution of rESCs in the whole body (B) and the forebrain (C) of the *Hesx1^-/-^* +mESCs and *Hesx1^-/-^*+rESCs chimeras (n = 3 for each group). All values are presented as the mean ± s.e.m.. *p < 0.05, **p < 0.01, ***p < 0.001, unpaired t-tests. ns, not significant.

### Cell autonomous and non-cell autonomous effects in forebrain reconstituted rat-mouse chimeras

We studied the brain development pace in WT (mouse), *Hesx1^-/-^* (mouse), *Hesx1*^-/-^+mESCs (mouse-mouse chimera), *Hesx1*^-/-^+rESCs (rat-mouse chimera) and WT (rat) embryos at E9.5 and E11.5 (n = 3 for each group) (Figure 6A). H&E-staining identified prosencephalon (PRO), mesencephalon (MS), rhombencephalon (RHO) in E9.5 mouse and E11.5 rat embryos, reflecting a ∼2 days delay in developmental timing in rats. At E11.5, mouse PRO expands into the telencephalon (T) and the diencephalon (D), the MS remains unchanged, and the RHO becomes the metencephalon (MT) and myelencephalon (MY)^39^. Intriguingly, *Hesx1*^-/-^+rESCs embryos exhibited the same pace of brain development to WT and *Hesx1*^-/-^+mESCs mice. These results suggest, similar to body and organ size, development of the rat brain tissues in rat-mouse chimeras is synchronized with the mouse host.

**Figure 6.**
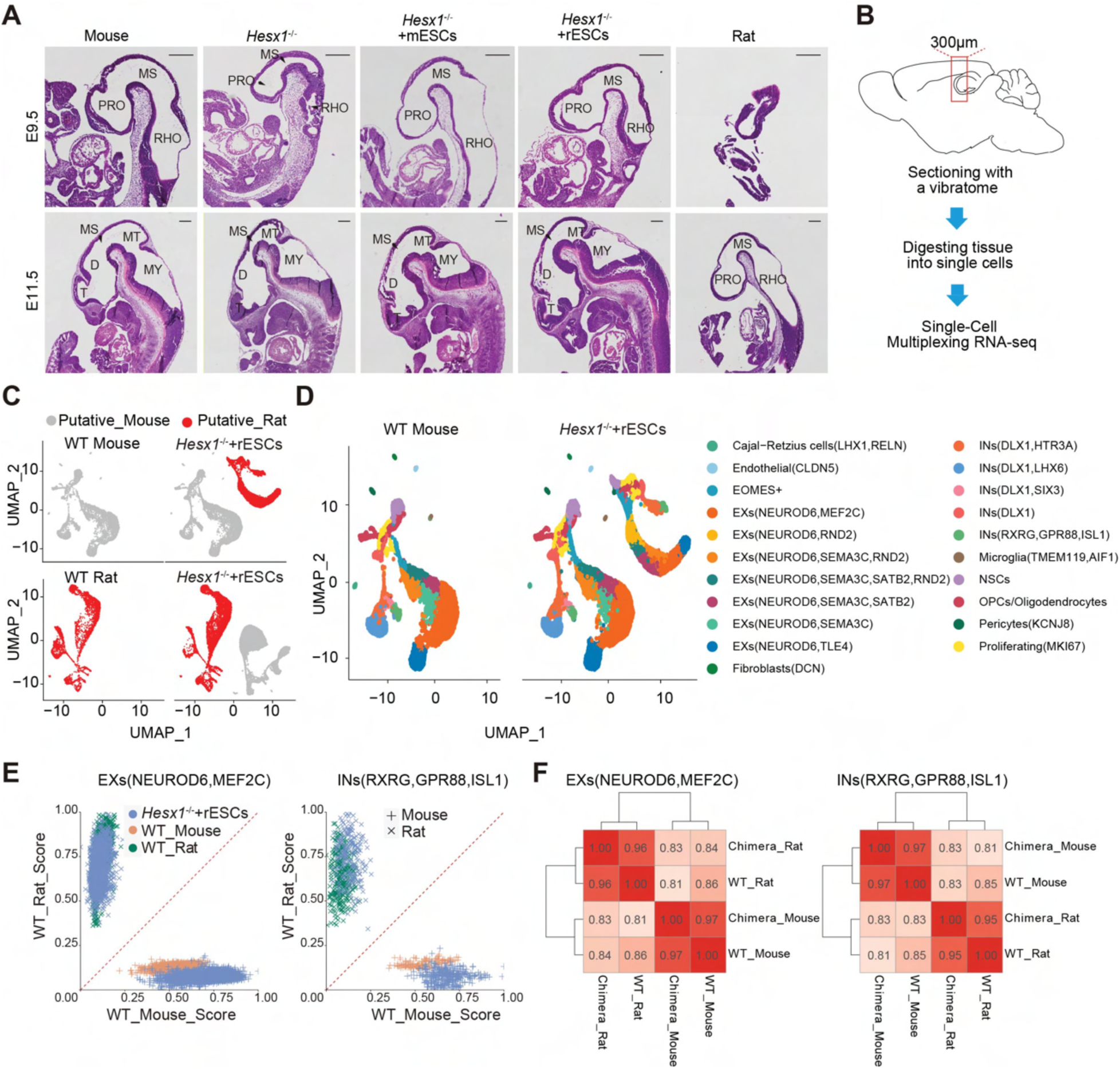
Cell autonomous and non-cell autonomous effects in forebrain reconstituted rat-mouse chimeras. (A) Representative H&E staining images of WT (mouse), *Hesx1^-/-^*(mouse), *Hesx1^-/-^*+mESCs (chimera), *Hesx1^-/-^*+rESCs (chimera) and WT (rat) embryos at E9.5 and E11.5 (n = 3 for each group). prosencephalon (PRO), mesencephalon (MS), rhombencephalon (RHO), telencephalon (T), diencephalon (D), metencephalon (MT) and myelencephalon (MY). Scale bars, 300 μm. (B) Schematic overview of single-cell RNA-seq analysis of forebrains tissues from *Hesx1*^-/-^ +rESCs chimeras (n = 3), WT rat (n = 1), and WT mouse (n = 1) at P0. (C) Uniform manifold approximation and projection (UMAP) visualization of single cells that were analyzed based on both their gene expression and species origin. Homologous genes between mice and rats were used for the analysis. Single cell data from WT mice (top) or rats (bottom) was used to perform integrative analysis for identifying rat cells and mouse cells in the *Hesx1^-/-^*+rESCs chimeras, respectively. Single cells from rats were highlighted as red dots. (D) Visualization of UMAP showing integrated analysis of 28,220 single cells between the WT mouse (left) and *Hesx1*^-/-^+rESCs chimeras (right). Cells are as color-coded by cell type. EXs, excitatory neuron; INs, inhibitory neurons; OPCs, oligodendrocyte progenitor cells; NSCs, neural stem cells. (E) Similarity score across cell types in *Hesx1^-/-^*+rESCs chimeras. Cells from WT mouse, WT rat, and chimeras are as color-coded. The putative species origins are as shape coded. (F) Heatmaps showing Pearson correlations between cells of the indicated sources. Pearson correlation was calculated based on the normalized gene expression levels. All values are presented as the mean ± s.e.m.. *p < 0.05, Unpaired t-test. ns, not significant.

To study the transcriptomic states of rESC-derived neurons, we isolated single cells using vibratome sections prepared from the cortex and hippocampus of *Hesx1*^-/-^+rESCs chimeras, WT mouse and WT rat, and subjected them to single-cell RNA-seq (scRNA-seq) (Figure 6B). Feature-barcode matrices of mouse and rat were separated based on reference mouse and rat genome. For each cell, the number of counts from the part of mouse and rat matrices were calculated respectively. Cells with counts 10-fold more from the rat than the counts from mouse were defined as putative rat cells (Figure 6C). Then, we identified different neuronal subtypes based on the expression of known markers (Figure S5A and S5B). We conducted 40× read depth DNA sequencing analysis on the cortex and hippocampus cells isolated from the *Hesx1*^-/-^+rESCs chimeras, which confirmed that *Hesx1* was successfully disrupted in the mouse cells (Figure S5C). scRNA-seq analysis revealed that rESCs generated various types of neurons in the chimeric forebrain including excitatory neurons, inhibitory neurons, glial cells, and others (n = 3 for *Hesx1*^-/-^+rESCs chimeras, n = 1 for WT mouse and WT rat) (Figure 6D and Figure S5D).

We performed comparative transcriptomic analysis of neurons from rat-mouse chimeras, as well as from WT rat and mouse. Our analysis revealed that rESC-derived neurons in rat-mouse chimeras transcriptomically resembled WT rat neurons while the expression profiles of host *Hesx1^-/-^* mouse neurons were similar to WT mouse neurons, suggesting cell autonomous rather than non-cell autonomous mechanisms dictate overall neuronal transcriptomic signatures in rat-mouse chimeras (Figures 6E and 6F; Figures S6A and S6B). Taken together, our findings demonstrate non-cell autonomous mechanisms determine the size and development time while cell autonomous mechanism shapes the transcriptomic landscape of rat forebrain tissues in mice generated via interspecies blastocyst complementation.

## DISCUSSION

In conclusion, we developed a rapid and efficient C-CRISPR-based blastocyst complementation (CCBC) platform and found that Hesx1 knockout in mouse blastocysts supported forebrain tissue generation from both mESCs and rESCs. It is important to note that while Dkk1 knockout supported the reconstitution of a functional forebrain derived from mESCs, it did not have the same effect with rESCs. Dkk1-null mice displayed agenesis phenotypes in both the forebrain and midbrain, suggesting that rESCs could not compensate for the broader developmental defects present in Dkk1-/- embryos compared to Hesx1-/- embryos. These findings emphasize the importance of screening and verifying the suitability of candidate genes for generating organ-complemented interspecies chimeras.

We anticipate that the C-CRISPR-based blastocyst complementation (CCBC) platform can be broadly applied to a wide range of organs, paving the way for utilizing large animals as hosts in blastocyst complementation experiments involving human cells. This approach has the potential to greatly expand our understanding of organ development, regeneration, and diseases, as well as address the global shortage of donor organs for transplantation. Despite the promising potential of CCBC, our study revealed that the chimeric rate of rESC-derived cells in mice gradually decreased as development progressed. This suggests the existence of additional xenogeneic barriers during mid-to-late prenatal development, which could impede the successful generation of chimeric organisms. Consequently, developing effective strategies to improve chimeric rates at mid-to-late gestation stages will be key to unlocking the full potential of CCBC.

Our study demonstrated that rESC-derived neurons are capable of not only functionally integrating but also enriching within a mouse forebrain. This discovery represents a crucial first step in realizing the full potential of interspecies neural blastocyst complementation as a transformative approach for studying brain development and disorders. The development of donor-derived brain tissues is heavily influenced by the host environment. However, the global gene expression profile strikingly maintains donor-specific characteristics, allowing researchers to investigate the complex interplay between extrinsic factors, such as the host environment, and intrinsic factors, such as species-specific gene expression. This dynamic interaction is essential in shaping the development of species-specific neuronal circuits and functions, and understanding these relationships can provide valuable insights into the underlying mechanisms of brain development, organization, and function.

With further advancements in interspecies chimerism, we anticipate that interspecies neural blastocyst complementation will pave the way for new opportunities to explore gene regulatory networks, cell-cell communication, and brain functions and behaviors within an evolutionary context.

## ACKNOWLEDGMENTS

We thank Dr. Mu-ming Poo for discussions and comments on the manuscript, and Optical Imaging facility Y. Wang, Y. Zhang and Q. Hu, and FACS facility S. Qian, H. Wu and L. Quan in ION. We also thank J. Du, M. Zhang, Z. Zhou, L. Zhang and Y. Zhao for technical assistance. This work was supported by NYSCF, NIH (HD103627-01A1), Welch (854671), R&D Program of China (2017YFC1001300 and 2018YFC2000100), CAS Strategic Priority Research Program (XDB32060000), National Natural Science Foundation of China (31871502, 31925016, 91957122, 31901047), Basic Frontier Scientific Research Program of Chinese Academy of Sciences. From 0 to 1 original innovation project (ZDBS-LY-SM001), Shanghai Municipal Science and Technology Major Project (2018SHZDZX05), Shanghai City Committee of Science and Technology Project (18411953700, 18JC1410100, 19XD1424400, 19YF1455100), and International Partnership Program of Chinese Academy of Sciences (153D31KYSB20170059). J.W. is a New York Stem Cell Foundation (NYSCF)–Robertson Investigator and Virginia Murchison Linthicum Scholar in Medical Research.

## AUTHOR CONTRIBUTIONS

J.H., H.Y. and J.W. conceptualized the project. H.Y., H.Z., F.G. and J.W. supervised the project. J.H. and B.H. designed experiments, derived and labeled the mESCs and rESCs. X.Y. and L.S. performed in vitro transcription and genotyping. J.H., X.Y. and Y.W. performed Microinjection. W.Y. performed mouse embryo transfer. J.H., Y.W. and S.L. performed behavior test. H.Z. performed stereotactic AAV injection. B.H., Y.F., Y.G. and Z.W. performed immunofluorescence analysis. X.L., Y.Z. and L.W. performed data analysis. J.H., C.L. and Y.A. performed single cell RNA sequencing experiments. J.H., H.Z., H.Y. and J.W. wrote the manuscript.

## METHODs

### Derivation of mESCs and rESCs

Mouse blastocysts were obtained from in vitro cultured mouse zygotes. The zona pellucida was removed with acid Tyrodes solution (Sigma), each blastocyst was transferred into a 4-well plate and cultured on the derivation medium: KnockOut DMEM (Gibco) supplemented with 20% KnockOut Serum Replacement (KSR, Gibco), 0.1mM non-essential amino acids (NEAA, Millipore), 1% penicillin/streptomycin (Gibco), 1% nucleosides (Millipore), 1% L-Glutamine (Gibco), 0.1mM β-mercaptoethanol (Millipore), 3 µM CHIR99021 (Selleck), 1 µM PD035901 (Selleck) and 1500 U/ml of mouse LIF (Millipore). After 5-7 days, the outgrowths of blastocysts were disaggregated and replated in the 2il medium. RESCs were derived as reported previously.^40^ Rat blastocysts were gently flushed out from the E4.5 timed-pregnant rats’ uteruses using M2 medium (Millipore). Zona pellucida was removed with acid Tyrodes solution (Sigma), and each blastocyst was transferred into a 4-well plate and cultured on MEFs with 2il medium or N2B27 plus 2il. After 5-7 days, the outgrowths of blastocysts were disaggregated and replated in the same culture medium.

### Cell culture

C57BL/6 ES and C57BL/6-tdTomato^+^ ES cell lines were cultured on feeder cells with 2il medium: DMEM (Millipore) supplemented with 15% FBS (Gibco), 0.1mM non-essential amino acids (NEAA, Millipore), 1% penicillin/streptomycin (Gibco), 1% nucleosides (Millipore), 1% L-Glutamine (Gibco), 0.1mM β-mercaptoethanol (Millipore), 3 µM CHIR99021 (Selleck), 1 µM PD035901 (Selleck) and 1000 U/ml of mouse LIF (Millipore). Rat ES cells derived from SD rats’ blastocyst were cultured on feeder cells with N2B27 plus 2il: Mix DMEM/F12 (Gibco) with Neurobasal medium (Gibco) at a ratio of 1:1, and then supplemented with 0.5% N-2 Supplement (Gibco), 1% B-27 Supplement (Gibco), 1% L-Glutamine (Gibco), 0.1mM β-mercaptoethanol (Millipore), 1 µM CHIR99021 (Selleck), 1 µM PD035901 (Selleck) and 10000 U/ml of recombinant rat LIF (ESGRO). All the cells were cultured in a 37℃ incubator under 5% CO_2_.

### Generation of tdTomato-labelled mouse and rESCs

C57BL/6 ES cells and SD rat ES cells were passaged at a ratio of 1:4-1:6 to a 6-well plate when cells reached 70% confluence. 5 µg plasmids (PBL-CAG-tdTomato-PBR: CMV-PBase = 1:1) were introduced into cells with Lipofectamine 3000 (Thermo Fisher Scientific) using the standard protocol. Cells were sorted out by BD FACS Aria II and cultured for blastocysts injection.

### In vitro transcription of Cas9 mRNA and sgRNAs

The T7 promoter sequence was added to the Cas9 coding region by PCR amplification of px260 (Addgene, 42229) using the primer pair listed in Table S3. The T7-Cas9 PCR product was purified using Gel Extraction Kit (Omega) and then used as the template for in vitro transcription (IVT) of Cas9 mRNA using the mMESSAGE mMACHINE T7 kit (Life Technologies). For sgRNA preparation, we added the T7 promoter sequence to the sgRNA template by PCR amplification of px330 (Addgene, 42230) using the primer pair listed in Table S3. The T7-sgRNA PCR product was purified using Gel Extraction Kit (Omega) and used as the template for IVT of sgRNAs using the MEGAshortscript T7 kit (Life Technologies). Both the Cas9 mRNA and sgRNAs were purified using the MEGAclear kit (Life Technologies) and eluted with elution buffer according to the standard protocol.

### Embryo and mice genotyping

Each zygote was collected into a PCR tube for genotyping when it reached to 4-cell stage. Zygotes were dissociated with 3 µl lysis buffer consists of 0.1% triton X-100 (Amresco, H2Q0212), 0.1% tween-20 (Amresco, H2T0301) and 30% proteinase K (20 min at 55℃ and then 5min at 95℃). For mice, toes were collected and genomic DNA was extracted for genotyping. All the primers used in this study were listed in Table S3.

### Microinjection of mRNAs into zygotes and establishment of blastocysts

For zygote preparation, fertilized eggs were obtained from super ovulated 8-week-old B6D2F1 female mice by intraperitoneal injection of 7.5 IU pregnant mare serum gonadotropin (PMSG, Ningbo Sansheng Medicine) followed by 7.5 IU human chorionic gonadotropin (hCG, Ningbo Sansheng Medicine) 48 hours later. After injection, the mice were mated with B6D2F1 males. One-cell-stage embryos were collected 22-24 hours after hCG injection. For zygote injection, the mixture of Cas9 mRNA (80 ng/μl) and sgRNA (25 ng/μl) was injected into the cytoplasm of fertilized embryos with well recognized two pronuclei in a droplet of M2 medium containing 5 μg/ml cytochalasin B (CB, Sigma) using a FemtoJet microinjector (Eppendorf) with continuous flow settings. Embryos were then cultured in KSOM+AA with D-Glucose (Millipore) at 37℃ under 5% CO_2_ for blastocyst injection. For blastocyst injection, blastocysts and ESCs were transferred into M2 medium droplets respectively. 8 to 12 mESCs or rESCs were introduced into the blastocoele near the ICM. The embryos were immediately transferred to recipients after injection.

### Transfer of mouse embryos

ICR female mice (8 weeks) in the stage of estrus were selected as recipients and mated with vasectomized ICR males overnight to induce pseudopregnancy. Injected E3.5 blastocysts were loaded into the embryo manipulation pipette and transferred into the uterine cavity of 2.5 dpc pseudo pregnant recipient. Each recipient received 15 to 20 blastocysts and this experiment was finished within 20 to 30 minutes.

### Whole genome sequencing and analysis for off-target effect

Genomes from *Hesx1^-/-^* (n = 6) and WT mice (n = 2) at P0 were sequenced with average 30 folds data using 150bp paired-end Illumina Xten platform. SolexaQA (V3.1.7.1)^41^ was used to filter the low-quality reads and Bwa-mem (0.7.16a)^42^ was used to align the clean reads to mm10 reference genome. Variant calling and filtration were performed using GATK (4.0.12.0)^42^ following GATK best practices. The parameters for variant filtration were “QD ‖ 2.0 || MQ < 40.0 || FS > 60.0 || SOR > 3.0 || MQRankSum < -12.5 || ReadPosRankSum < -8.0”. Cas-offinder^43^ was used to predict the off-target sites with no more than five mismatches. Overlapping of SNVs or Indels was performed using Bedtools(v2.29.2).^44^ Dbsnp database and UCSC repeat tracks were used to filter the variants. All of the off-target sites were further examined by PCR amplification and sequencing, and none of true off-target sites were found (Table S2).

### Immunofluorescence analysis

Mice were anaesthetized and transcardially perfused with normal saline and 4% paraformaldehyde at P0, P7 or adult stage. Brain and other tissues or organs were removed and fixed with 4% paraformaldehyde at 4℃ for 24h, dehydrated with 30% sucrose in 0.1 M sodium phosphate buffer overnight at 4 °C, and then embedded in OCT. Brain and pancreas slices were sectioned into slices with thickness of 30 µm, washed with PBS three times, and then incubated with primary antibody at 4°C overnight. The following primary antibodies were used: anti-RFP (rabbit polyclonal, 1:500; 600401379, Rockland), Ctip2 (rat monoclonal, 1:200; ab18465, Abcam), Alpha Amylase (rabbit polyclonal, 1:200; PA5-51078, Thermo Fisher Scientific), Insulin (rabbit polyclonal, 1:1000; 15848-1-AP, Proteintech), Glucagon (rabbit polyclonal, 1:1000; 15954-1-AP, Proteintech). Sections were washed three times with PBS and incubated with secondary antibody Alexa Fluor 488-AffiniPure Donkey Anti-Rabbit IgG (H+L) (1:500, 711545152, Jackson), Alexa Fluor 488-AffiniPure Donkey Anti-Rat IgG (H+L) (1:500, 712545153, Jackson) and Cy^TM^3-AffiniPure Goat Anti-Rabbit IgG (H+L) (1:500, 111165003, Jackson) for two hours at room temperature. Sections were washed three times with PBS, mounted with Vectashield (Vector Laboratories), a mounting medium containing DAPI (Thermo Fisher Scientific) for nuclear counterstaining. Non-brain tissues (liver, heart, stomach,spleen, kidney, tail and eye) were fixed and sectioned similarly to the brains. Slides were imaged with vs120 and FV3000 (Olympus) and processed using Image J.

### Measurement of the width of the hippocampus and the cortex

To measure the layer width of the cortex and width of the hippocampus, brain slices from P7, adult chimeras (*Dkk1*^-/-^+mESCs, *Hesx1*^-/-^+mESCs, and *Hesx1*^-/-^+rESCs) and conventional chimera controls (WT+mESCs and WT+rESCs) were analyzed. Width of layer I–VI, I, II–IV and VI were determined by DAPI staining and the boundary of cortex. Layer V was determined by Ctip2 immunostaining. Width of CA1, CA3 and dentate gyrus were determined by DAPI staining. For quantification, target brain regions with high structural similarity were selected (at least three sections per mouse).

### Analysis of single-cell RNA-seq data

For the raw sequencing data, the Cell Ranger (v3.1; 10X Genomics) was used for reads alignment, deduplication and quantification of gene expression with default parameters. First, both the Mouse (Mus musculus, GRCm38.p6) and Rat (Rattus norvegicus, Rnor_6.0) reference genomes were used for reads alignment, following the instructions from the 10*×* Genomics. Raw reads were aligned to the two-species reference genome and quality checked. Then, those uniquely mapped reads were used for UMI counting. Finally, for each barcode observed, gene expression was quantified and those cells with much lower RNA content were discarded. The filtered feature-barcode matrices were constructed by the Cell Ranger (v3.1; 10X Genomics).

The cell source in the chimera was first identified. Since the two-species reference genome was used, feature-barcode matrices could be separated into the part of Mouse and Rat, respectively. For each cell, the number of counts from the part of Mouse and Rat was calculated, respectively. Cells with counts 10-fold more from the Mouse than the counts from Rat were considered the Mouse cells. And cells with counts 10-fold more from the Rat than the counts from Mouse were considered the Rat cells. The remaining cells were considered as the ambiguous and were excluded. As quality control, cells were included if they passed the following filters: 1) the percentage of counts from the mitochondrial genome was less than 10%, 2) the number of unique detected counts was more than the half and less than the two-fold of the average unique detected counts across the cells within the same sample. For chimera, quality control was performed based on the source of cell, 3) the ratio of log10-transformed number of detected genes and log10-transformed number of detected counts was higher than 0.85. In total, 35,205 out of 58,688 cells were used for the downstream integrated analysis by Seurat (v3.2.3),^45^ according to the recommended procedures.^45^

For identifying cell types, we defined a set of marker genes for each cell type based on previously similar studies.^45–52^ For Cajal-Retzius cells: *LHX1*, *RELN*. For endothelial: *CLDN5*. For fibroblasts: *DCN*. For Microglia: *TMEM119*, *AIF1*. For neural stem cells (NSCs): *GFAP*. For proliferating: *CENPF*, *MKI67*, *TOP2A*. For Oligodendrocyte precursor cells (OPC) / Oligodendrocyte: *OLIG1*, *OLIG2*, *PDGFRA*. For pericytes: *KCNJ8*. For subgroups of excitatory neuron (Ex): *EOMES*, *NEUROD6*, *MEF2C*, *RND2*, *SEMA3C*, *SATB2*, *TLE4*. For subgroups of inhibitory neuron (In): *DLX1*, *HTR3A*, *LHX6*, *SIX3*, *RXRG*, *GPR88*, *ISL1*.

To projection cells onto different brain regions, markers for brain regions were retrieved^53^. For each cell, expression score was calculated for each region and the region with highest score was assigned to the cell.

### Selection of gene set for comparative species analysis

List of homologous genes between Mouse (Mus musculus) and Rat (Rattus norvegicus) was downloaded from MGI Vertebrate Homology (http://www.informatics.jax.org/downloads/reports/HOM_AllOrganism.rpt). Those homologous genes between mouse and rat were used for the comparative species analysis as previously described.^46^

### 40 × read depth DNA Sequencing

The PCR products amplified from Mouse *Hesx1* genomic sites of *Hesx1*^-/-^+rESCs chimeras were sequenced with average 40 folds data using 150bp paired-end Illumina Xten platform. Barcodes for demultiplexing:

*Hesx1*^-/-^+rESCs1#^GAATTCATCACGGGATCCACATATAAAATACTGCCACT;
*Hesx1*^-/-^+rESCs2#^GAATTCCGATGTGGATCCACATATAAAATACTGCCACT;
*Hesx1*^-/-^+rESCs3#^GAATTCTTAGGCGGATCCACATATAAAATACTGCCACT.

Sequencing reads were demultiplexed using cutadapt (v2.10).^54^ Alignment of amplicon sequences to a reference sequence was performed using CRISPResso2.^55^ For the quantification of editing efficiency, editing efficiency was calculated as: percentage of (number of modified reads) / (number of total reads). CRISPResso2 was run with default parameters except the ‘exclude_bp_from_left’ was set to zero.

### Morris water maze (MWM)

In the preparation stage, the three lights on the wall of behavior laboratory were turned on, the escape platform remained in the center of one quadrant of the pool. The pool was filled with water that has a temperature of 20-23℃. To hide the platform, the platform was submerged below the water surface (0.5-1 cm) and soluble non-toxic white powder was added into the water. For learning stage (day 1 to 5), adult mice were placed in the pool facing the tank wall, each mouse was given 60s to swim to find the hidden platform. The mouse was allowed to stay there for 2s if it could find the platform. If the mice could not find the platform within 60s, they were transferred from the water to the platform. Each mouse completed four learning trails on day 1 to day 5, and each trail started from different orientations of the pool. To assess the spatial learning and memory, the platform was removed in the testing stage (day 6) and mice were placed in the quadrant opposite to the quadrant where the platform was located previously and given 60s to swim. Time spent and path length in each quadrant were recorded. The swim trajectory was recorded using water maze 5 software.

### Contextual fear conditioning

Contextual fear conditioning was done as previously describe^34^. Mice were placed in a system comprising four boxes that have iron cages and different contexts. All animals completed 2 trails (5 minutes per trial) each day and this task was repeated on three consecutive days. On day one, mice were placed in context A that was used for fear conditioning. After 180s, the mice were received the first foot shock (0.75mA, 180s), and the second foot shock was added 30s later. On day two, mice were placed in the context A again for 5 mins. On day three, mice were placed in context B for 5 mins. The boxes were cleaned with 70% ethanol before placing the mice. The movements were detected by the infrared sensors and the freezing behavior was recorded using FreezeFrame (4.07) software.

### Open-field assay

The open-field assay took place in a square, white Plexiglas box. Mice were placed in the arena and allowed to move freelyfor about 30 minutes while being recorded and analyzed by EthoVisonXT 11.5 for the following parameters: distance moved, velocity, and time spent in pre-defined zones.

### AAV injection

The AAVs (serotype: AAV8) were injected into the ALM, similar to previously reported^32^. In brief, mice were anaesthetized with isoflurane (R510-22, RWD) and the head was fixed on the stereotactic frame. After cutting the skin over the skull, a micropipette was advanced into the ALM (AP +2.3 mm, ML ±1.7 mm and DV −0.6 mm) using a micromanipulator and 0.1-0.2 μl AAV (AAV8-hSyn-EGFP-WPRE-pA, titer: 1 **×** 10^13^ vg/ml) in the micropipette were released. Two weeks later, brains were collected and sectioned for immunostaining and analysis.

### Quantification and statistical analysis

All values are shown as mean ± s.e.m.. Unpaired Student’s t-tests (two tailed) were used to evaluate statistical significance (*p < 0.05, **p < 0.01, ***p < 0.001). Randomization was used in all experiments.

## Supplemental Figures

**Figure S1.**
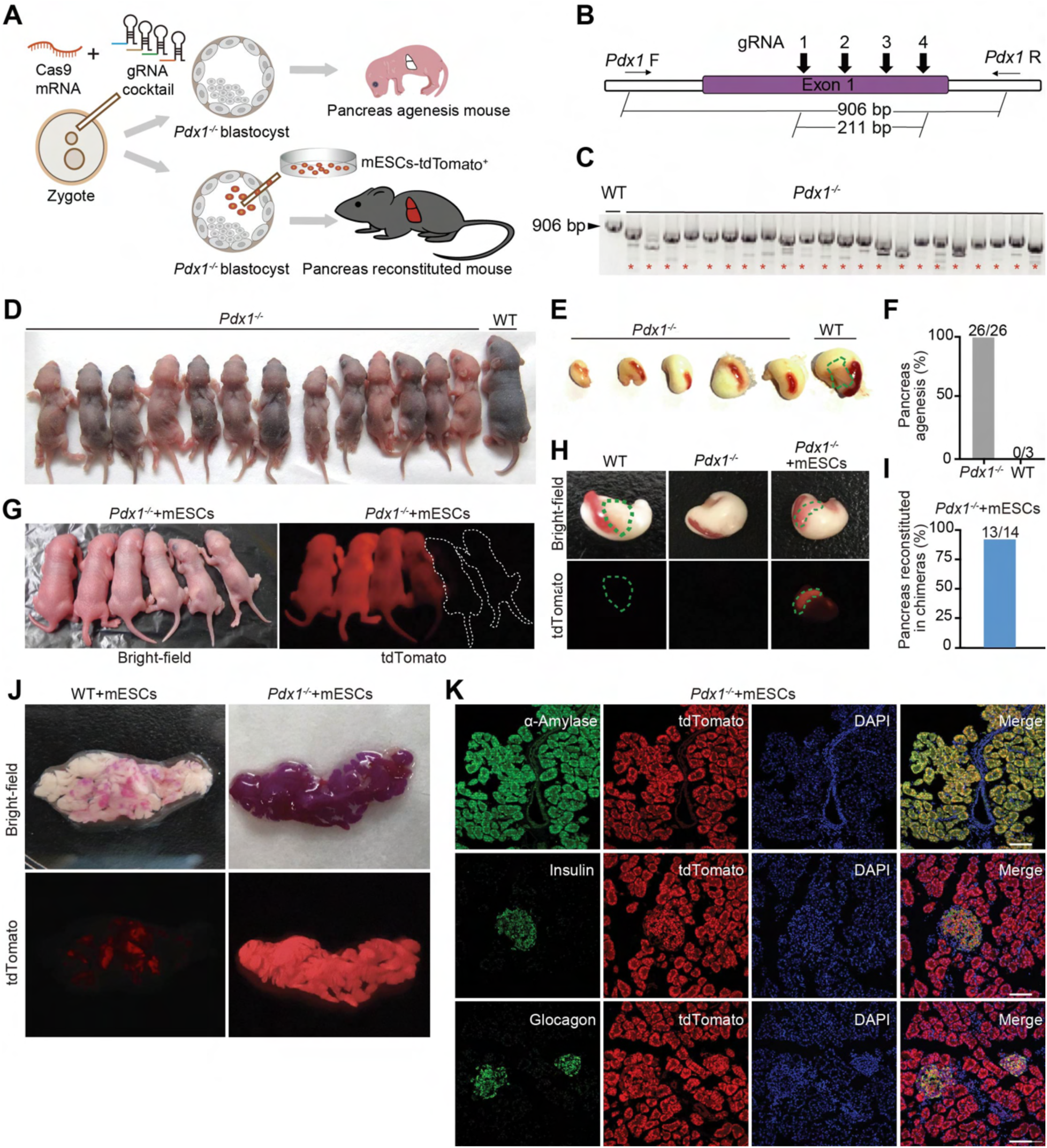
CCBC enables one-step generation of donor mESC-derived pancreatic tissue in *Pdx1^-/-^* mice. (A) Schematic showing generation of pancreas reconstituted mice using CCBC. *Cas9* mRNA and 4 sgRNAs targeting *Pdx1* were co-injected into WT mouse (B6D2F1) zygotes. The compensated blastocysts could develop into normal individuals with an organ derived from tdTomato^+^ mESCs or rESCs. (B) Design of 4 sgRNAs targeting *Pdx1*. Note that the PCR products of knockout blastocysts are about 200 bp shorter than that of WT. (C) PCR-based genotyping of 4-cell embryos. Red asterisks indicate the *Pdx1* knockout embryos. (D) Photograph of newborn *Pdx1^-/-^* and WT mice at postnatal day (P) 2. (E) Macroscopic images of stomach, spleen and pancreas in *Pdx1^-/-^* and WT mice at P2. (F) Quantification of the efficiency of pancreas agenesis, showing that injection of *Cas9* mRNA and 4 sgRNAs lead to the pancreas agenesis of all mice. Numbers above the bar indicates the number of mice. (G) Representative images showing *Pdx1^-/-^*+mESCs mice (*Pdx1^-/-^*+mESCs, *Pdx1*^-/-^ mice with mESC-derived pancreas) at P1. Note that the 4 mice on the left were pancreas-complemented, and the 2 mice on the right failed to develop a tdTomato^+^ mESC-derived pancreas. (H) Representative images showing the reconstitution of pancreas in *Pdx1^-/-^*+mESCs chimeras at P1. Note that the WT mouse has a normal pancreas, the *Pdx1^-/-^* mouse has no pancreas and the *Pdx1^-/-^*+mESCs chimera has a tdTomato labeled normal pancreas. (I) Quantification of pancreas reconstitution efficiency in the *Pdx1^-/-^*+mESCs chimeras (J) Macroscopic images showing the mESC-derived pancreases generated in *Pdx1^-/-^*+mESCs (n = 4 chimeras) and WT+mESCs (n = 3 chimeras) at 8 weeks of age. (K) Immunofluorescence images showing the contribution of mESC-derived cells in a reconstituted pancreas (n = 6 chimeras). Note that α-Amylase is a pancreatic exocrine marker; insulin and glucagon are pancreatic endocrine markers. Scale bar, 100 μm.

**Figure S2.**
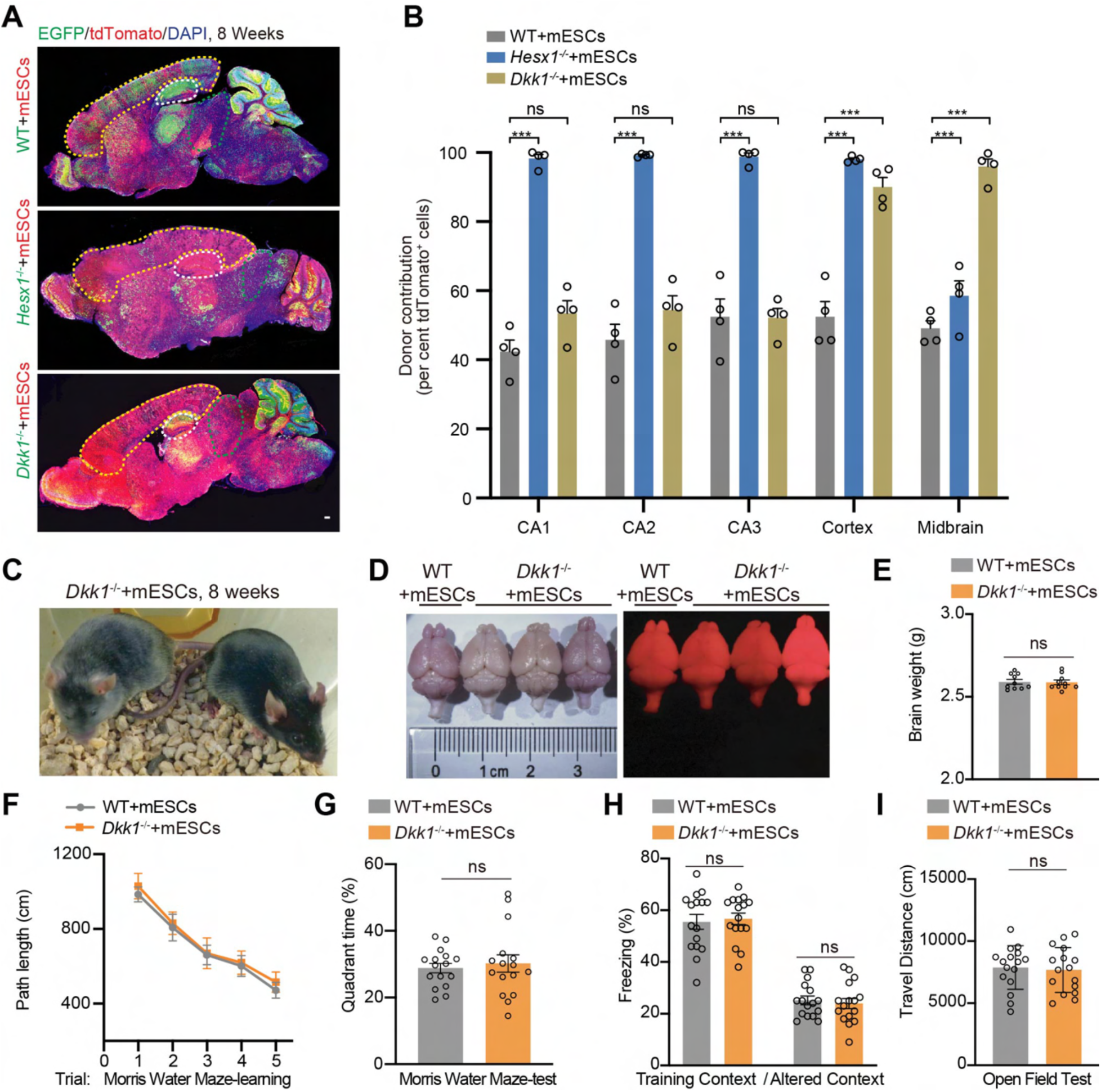
*Dkk1^-/-^*+mESCs chimeras are structurally and functionally normal. (A) Representative sagittal brain sections from WT+mESCs (upper, n = 4), *Hesx1^-/-^*+mESCs (middle, n = 4) and *Dkk1^-/-^*+mESCs (lower, n = 4) chimeras at 8 weeks of age. Yellow dashed line, cortex; white dashed line, hippocampus; green dashed line, midbrain. Scale bar, 200 μm. (B) Quantification of donor ES cell contribution in WT+mESCs, *Hesx1^-/-^*+mESCs and *Dkk1^-/-^* +mESCs chimeras at 8 weeks of age. n = 4 for each group. (C) Representative images of *Dkk1^-/-^*+mESCs chimeras (8 weeks of age). (D) Bright-field and fluorescence images showing no obvious difference between the brains of *Dkk1^-/-^*+mESCs and WT+mESCs chimeras (10 weeks of age). (E) Brain weight of *Dkk1^-/-^*+mESCs and WT+mESCs chimeras (n = 10, 10 weeks of age). (F) Mean path length to platform for WT+mESCs (gray) or *Dkk1^-/-^*+mESCs (orange) chimeras in learning trials. (G) Mean time ratio in the target quadrant for WT+mESCs (gray) and *Dkk1^-/-^*+mESCs (orange) chimeras in test trials. (H) Contextual fear conditioning test of WT+rESCs (gray) and *Dkk1^-/-^*+mESCs (orange) chimeras. (I) Open-field assay of WT+mESCs (gray) and *Dkk1^-/-^*+mESCs (orange) chimeras. For (F) to (I), n = 16 chimeras for each group. All values are presented as mean ± s.e.m.. *p < 0.05, **p < 0.01, ***p < 0.001, Unpaired t-test. ns, not significant.

**Figure S3.**
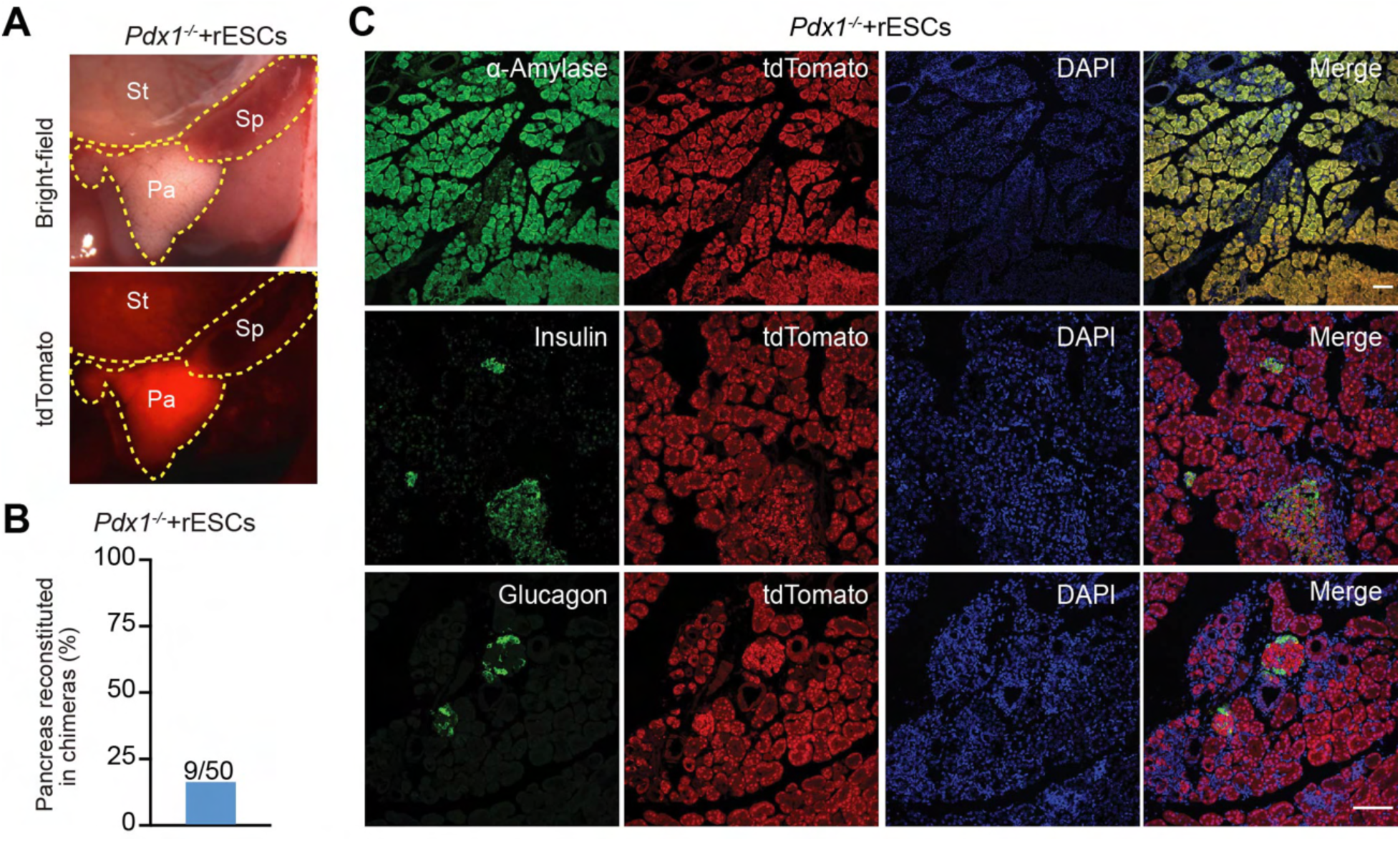
Generation of a rat pancreas in mouse via CCBC. (A) Macroscopic images showing a rESC-derived pancreas generated in *Pdx1^-/-^*+rESCs chimeras (n = 4) at 8 weeks of age. (B) Quantification of pancreas complementation efficiency in *Pdx1^-/-^*+rESCs chimeras. (C) Immunofluorescence images showing the distribution of rESC-derived cells in a reconstituted pancreas from *Pdx1^-/-^*+rESCs chimeras (n = 4 chimeras). Note that α-Amylase is a pancreatic exocrine marker, insulin and glucagon are pancreatic endocrine markers. Scale bar, 100 μm.

**Figure S4.**
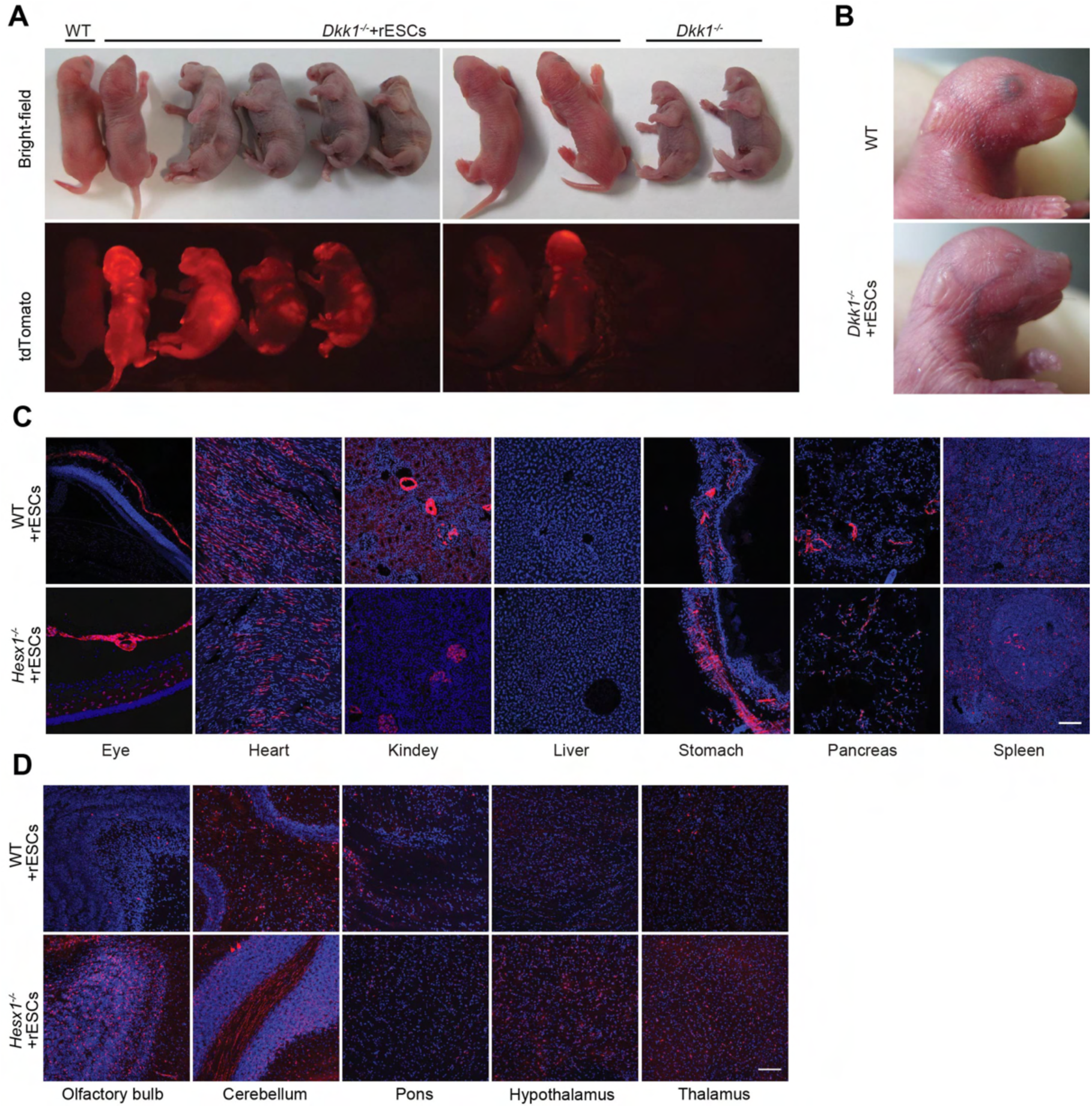
Chimeric contribution of rESCs in mice. (A) Bright-field and fluorescence images showing WT mice and *Dkk1^-/-^*+rESCs partial reconstituted chimeras at P0. (B) Bright-field images showing the head of a WT mice and a forebrain partial reconstituted *Dkk1^-/-^*+rESCs chimera. (C) Representative images showing the contribution of tdTomato^+^ donor rESCs to the indicated non-brain organs in WT+rESCs and *Hesx1^-/-^*+rESCs chimeras (n = 3, 10 weeks of age). Red, rESC-derived cells; blue, DAPI. Scale bar, 100 μm. (D) Representative images showing the contribution of tdTomato^+^ donor rESCs to different brain regions in WT+rESCs and *Hesx1^-/-^*+rESCs chimeras (n = 3 per group, 10 weeks of age).

**Figure S5.**
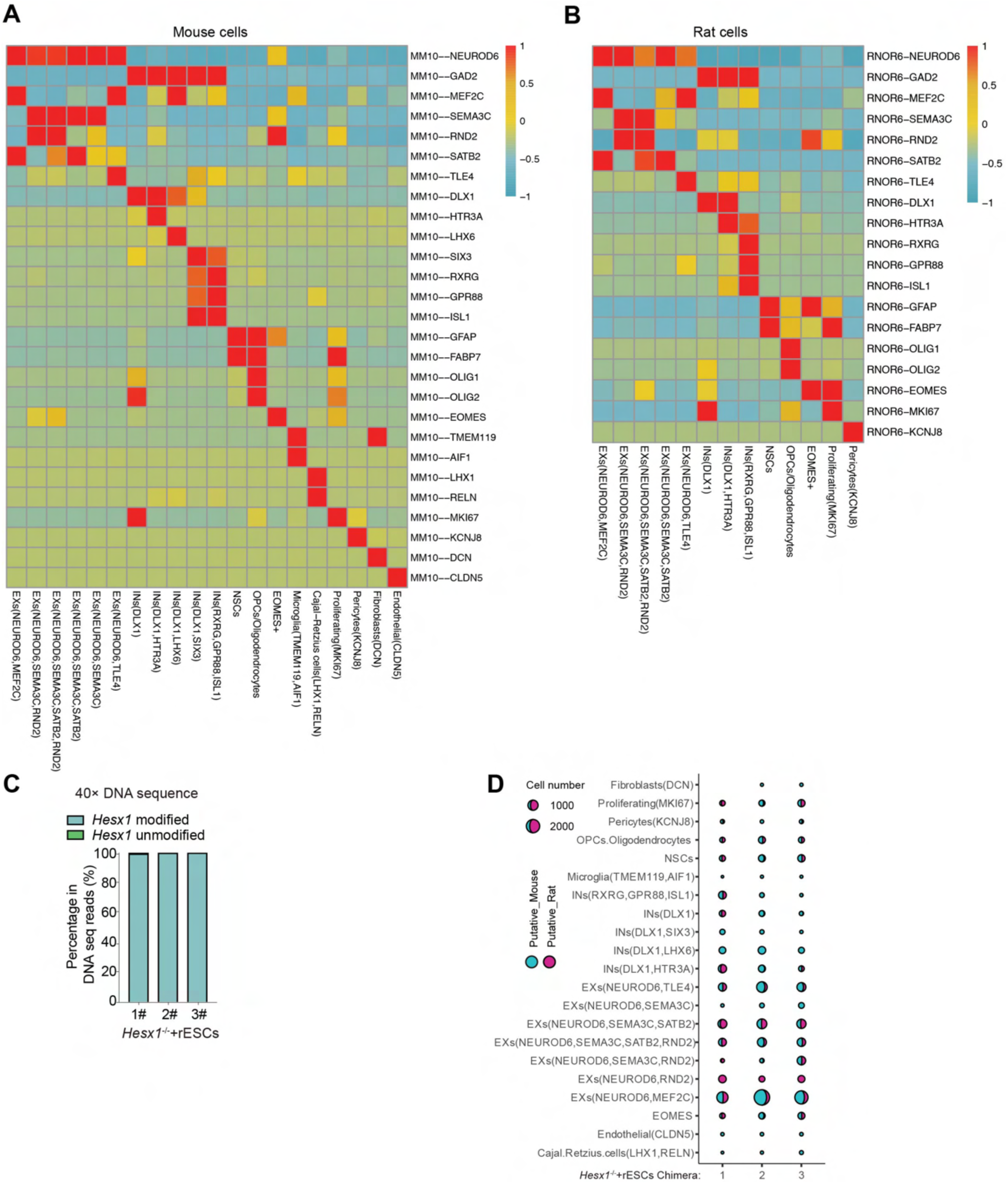
Integrated analysis for identifying cell types in *Hesx1^-/-^*+rESCs chimeras. (A) and (B) Heatmaps of selected cell-type-specific genes for the putative rat (left) and mouse cells (right) based on their expression level. INs, inhibitory neuron; EXs, excitatory neuron; OPCs, oligodendrocyte progenitor cells; NSC, neural stem cell. The average expression level of each gene was calculated and scaled using the Seurat (v3.1.4) R package. (C) Quantification of the editing efficiency via deep sequencing analysis. The editing efficiency was calculated as: percentage of (number of modified reads) / (number of total reads). *Hesx1* modified and unmodified reads were represented as blue and green, respectively. (D) Dot plot showing the proportional representation of putative mouse/rat cells across cell types identified in chimeras. The total number of cells within the cell type was encoded by the size of the dot, while the color encodes the putative source of cells.

**Figure S6.**
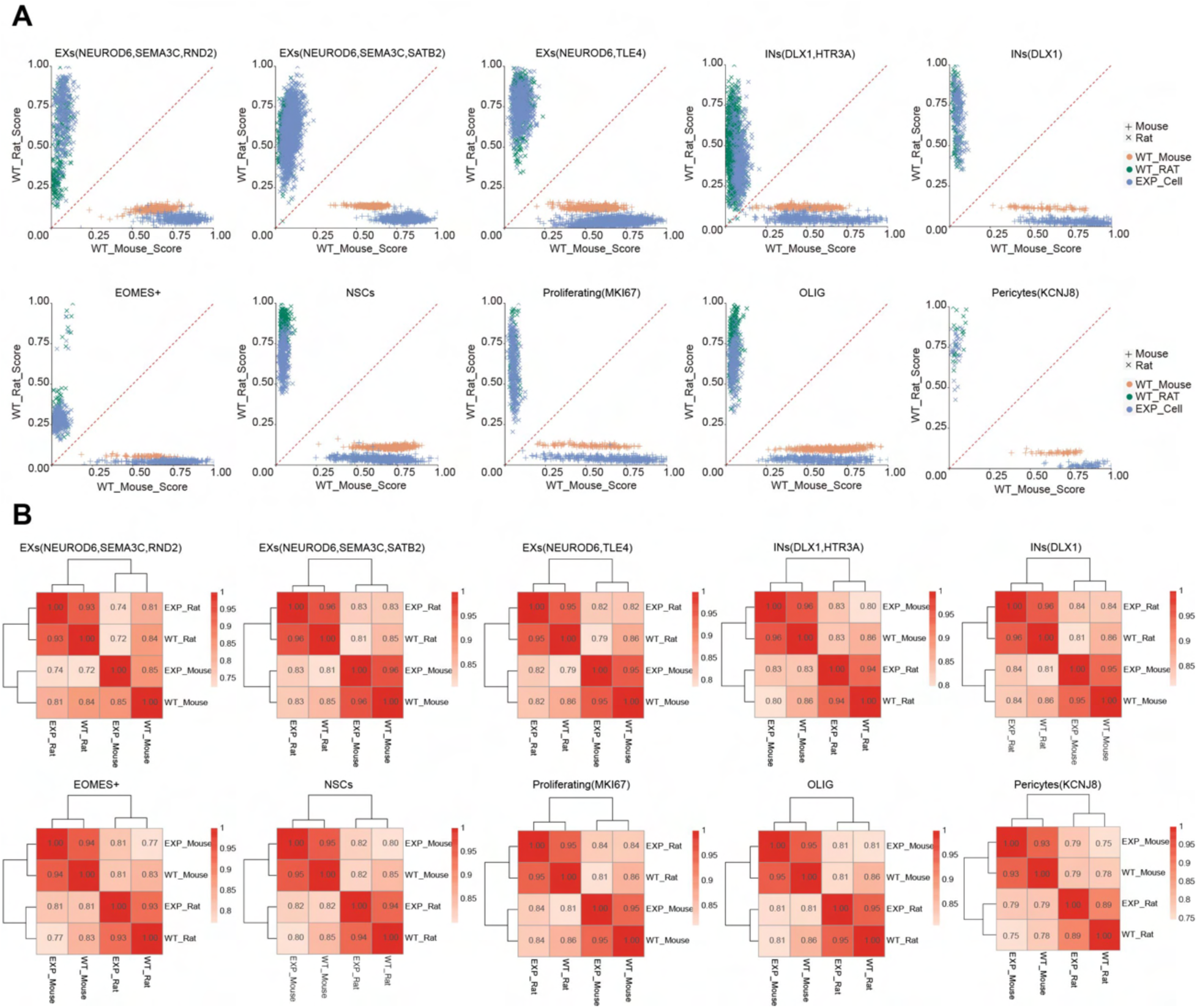
Similarity score and Pearson correlations across cell types in *Hesx1^-/-^*+rESCs chimeras. (A) Similarity score across cell types in *Hesx1^-/-^*+rESCs chimeras, WT mouse and WT rat are as color-coded. The putative source of cells is as shape coded. WT, wildtype. EXP, *Hesx1*^-/-^+rESCs. (B) Heatmaps showing Pearson correlations between cells of the indicated sources. Pearson correlation was calculated based on the normalized gene expression levels. WT, wildtype. EXP, *Hesx1*^-/-^+rESCs.

