## Supplementary material for "Interspecies blastocyst complementation generates functional rat cell-derived forebrain tissues in mice": Tables S1-S3

### Table S1. Summary of embryo transfer and newborn.

**B6D2F1 zygotes**

**Mouse-Mouse**

| Targeted organ | Gene konckout | Cas9/sgRNA (ng/μL) | ESCs injected | Transferred Embryos (Recipients) | Newborn pups | Chimeric pups | Reconstituted pups (% of chimeras) | Partial reconstituted pups (% of chimeras) | Agenesis pups (% of newborn) |
|---|---|---|---|---|---|---|---|---|---|
| Pancreas | *Pdx1* | 80/25*4 | / | 146(6) | 26 | / | / | / | 26(100%) |
| | | | mESCs | 159(7) | 37 | 14 | 13(93%) | 1(7%) | 23(62%) |
| Midbrain and forebrain | *Dkk1* | 80/25*4 | / | 122(5) | 24 | / | / | / | 24(100%) |
| | | | mESCs | 460(21) | 101 | 61 | 55(90%) | 6(10%) | 40(40%) |
| Forebrain | *Hesx1* | 80/25*4 | / | 51(3) | 11 | / | / | / | 11(100%) |
| | | | mESCs | 445(20) | 130 | 68 | 66(97%) | 2(3%) | 62(48%) |
| / | / | / | mESCs | 130(6) | 27 | 12 | / | / | / |

**Rat-Mouse**

| Targeted organ | Gene konckout | Cas9/sgRNA(ng/μL) | ESCs injected | Transferred Embryos (Recipients) | Newborn pups | Chimeric pups | Reconstituted pups (% of chimeras) | Partial reconstituted pups (% of chimeras) | Agenesis pups (% of newborn) |
|---|---|---|---|---|---|---|---|---|---|
| Pancreas | *Pdx1* | 80/25*4 | rESCs | 910(30) | 360 | 50 | 9(18%) | 41(82%) | 310(86%) |
| Forebrain | *Hesx1* | 80/25*4 | rESCs | 6836(271) | 2089 | 417 | 16(4%) | 401(96%) | 1672(80%) |
| Midbrain and forebrain | *Dkk1* | 80/25*4 | rESCs | 5416(216) | 1715 | 374 | / | 374(100%) | 1341(78%) |
| / | / | / | rESCs | 311(13) | 67 | 13 | / | / | / |

**C57BL/6 zygotes**

| Targeted organ | Gene konckout | Cas9/sgRNA(ng/μL) | ESCs injected | Transferred Embryos (Recipients) | Newborn pups | Chimeric pups | Reconstituted pups (% of chimeras) | Partial reconstituted pups (% of chimeras) | Agenesis pups (% of newborn) |
|---|---|---|---|---|---|---|---|---|---|
| Pancreas | *Pdx1* | 80/25*4 | mESCs | 48(3) | 13 | 7 | 7(100%) | 0 | 6(46%) |

Note: Reconstituted pups are defined as tdTomato positive-chimeras that target organ is intact, and partial reconstituted pups are defined as tdTomato positive-chimeras that targeted organ is agenesis.

### Table S2. Off-target analysis of *Hesx1* gRNAs.

#### Predicted off-target sites of 4 *Hesx1* gRNAs

| sgRNA | No. of Mismatches | No. of NGG genomic sites | Off-target sites 1# | 2# | 3# | 4# | 5# | 6# |
|---|---|---|---|---|---|---|---|---|
| *Hesx1* gRNA 1+2+3+4 | 0 | 4 | NA | NA | NA | NA | NA | NA |
| | 1 | 1 | 0 | 0 | 0 | 1 | 0 | 0 |
| | 2 | 4 | 0 | 0 | 0 | 1 | 0 | 0 |
| | 3 | 78 | 0 | 0 | 0 | 0 | 0 | 0 |
| | 4 | 1,100 | 0 | 0 | 0 | 0 | 0 | 0 |
| | 5 | 3,617 | 2 | 2 | 2 | 0 | 0 | 0 |

#### PCR identification of the off-target sites

| Mouse | sgRNA | Chromosome | Start | End | Location | Reference | Variation | Indel frequency | Off-target effect |
|---|---|---|---|---|---|---|---|---|---|
| 1# | 3 | Chr1 | 165672421 | 165672441 | 165672429 | G | GGGCGCAATGACGGTTTTACCATGCTCTGTGTGTGGGGCAGGC GCAATGACGGTTTTACCATGCTCTGTGTGTGGGGCAGGCGCAA TGACGGTTTTACCATGCTCTGTGTGGGGCAGGCGCAATGACGG TTTTACCATGCTCTGTGTGGGGCA | 1/9 | 0 |
| 1# | 3 | Chr13 | 66994058 | 66994078 | 66994073 | A | ACACACACACACACACACACACACACACACACACACAAC | ND | ND |
| 2# | 4 | Chr8 | 33656450 | 33656470 | 33656455 | CCTGGCTT | C | 0/8 | 0 |
| 2# | 4 | Chr8 | 34760311 | 34760331 | 34760313 | AG | A | 7/13 | 0 |
| 3# | 4 | Chr8 | 33656450 | 33656470 | 33656455 | CCTGGCTT | C | 0/7 | 0 |
| 3# | 4 | Chr8 | 34760311 | 34760331 | 34760313 | AG | A | 2/7 | 0 |
| 4# | 4 | Chr4 | 47696292 | 47696312 | 47696300 | ACAGAGTC | A | 0/9 | 0 |
| 4# | 4 | Chr11 | 114574549 | 114574569 | 114574566 | C | CA | 6/10 | 0 |
| 6# | 4 | Chr5 | 103299857 | 103299877 | 103299863 | C | CT | ND | ND |

### Table S3. Primers used in this study.

Primers used for generating PCR products to serve as substrates for T7 transcription.

| Name | Sequence (5'→3) |
|---|---|
| fwd_Cas9 mRNA_T7 | TAATACGACTCACTATAGGGAGATTTCAGGTTGGACCGGTG |
| rev_Cas9 mRNA_T7 | GACGTCAGCGTTCGAATTGC |
| rev_sgRNA_T7 | AAAAGCACCGACTCGGTGCC |
| fwd_mouse *Pdx1* gRNA 1 | TAATACGACTCACTATAGGG tcgtatggggagatgtccgg GTTTTAGAGCTAGAAATAG |
| fwd_mouse *Pdx1* gRNA 2 | TAATACGACTCACTATAGGG tgggcgagcccgagctgagc GTTTTAGAGCTAGAAATAG |
| fwd_mouse *Pdx1* gRNA 3 | TAATACGACTCACTATAGGG ggacgcggttgggctcttcc GTTTTAGAGCTAGAAATAG |
| fwd_mouse *Pdx1* gRNA 4 | TAATACGACTCACTATAGGG tcacgcgtggaaaggccagt GTTTTAGAGCTAGAAATAG |
| fwd_mouse *Pax6* gRNA 1 | TAATACGACTCACTATAGGG ctgggcaggtattacgagac GTTTTAGAGCTAGAAATAG |
| fwd_mouse *Pax6* gRNA 2 | TAATACGACTCACTATAGGG acttctggagtcgccactct GTTTTAGAGCTAGAAATAG |
| fwd_mouse *Pax6* gRNA 3 | TAATACGACTCACTATAGGG agggcactcccgtttatact GTTTTAGAGCTAGAAATAG |
| fwd_mouse *Pax6* gRNA 4 | TAATACGACTCACTATAGGG ccgagacagattattatccg GTTTTAGAGCTAGAAATAG |
| fwd_mouse *Lhx1* gRNA 1 | TAATACGACTCACTATAGGG cgcgggagggaagtactcac GTTTTAGAGCTAGAAATAG |
| fwd_mouse *Lhx1* gRNA 2 | TAATACGACTCACTATAGGG cgacagggcaattagagtcg GTTTTAGAGCTAGAAATAG |
| fwd_mouse *Lhx1* gRNA 3 | TAATACGACTCACTATAGGG gggctgcaaaaggcccatcc GTTTTAGAGCTAGAAATAG |
| fwd_mouse *Lhx1* gRNA 4 | TAATACGACTCACTATAGGG tgcatttacattcacagcac GTTTTAGAGCTAGAAATAG |
| fwd_mouse *Otx2* gRNA 1 | TAATACGACTCACTATAGGG ccaccccccggaaacagcga GTTTTAGAGCTAGAAATAG |
| fwd_mouse *Otx2* gRNA 2 | TAATACGACTCACTATAGGG tagggcacagctcgacgttc GTTTTAGAGCTAGAAATAG |
| fwd_mouse *Otx2* gRNA 3 | TAATACGACTCACTATAGGG gatgtctgggtaccgggtct GTTTTAGAGCTAGAAATAG |
| fwd_mouse *Otx2* gRNA 4 | TAATACGACTCACTATAGGG cctacctgcaccctggattc GTTTTAGAGCTAGAAATAG |
| fwd_mouse *DKK1* gRNA 1 | TAATACGACTCACTATAGGG gcgccgggagttctctatga GTTTTAGAGCTAGAAATAG |
| fwd_mouse *DKK1* gRNA 2 | TAATACGACTCACTATAGGG aagaacctgcccccaccgct GTTTTAGAGCTAGAAATAG |
| fwd_mouse *DKK1* gRNA 3 | TAATACGACTCACTATAGGG tttgcgtccttcggagatga GTTTTAGAGCTAGAAATAG |
| fwd_mouse *DKK1* gRNA 4 | TAATACGACTCACTATAGGG tggcactggctcctagcaga GTTTTAGAGCTAGAAATAG |
| fwd_mouse *Cer1* gRNA 1 | TAATACGACTCACTATAGGG ggcacggccacaaacagatc GTTTTAGAGCTAGAAATAG |
| fwd_mouse *Cer1* gRNA 2 | TAATACGACTCACTATAGGG ggggggtaaaattcggtctc GTTTTAGAGCTAGAAATAG |
| fwd_mouse *Cer1* gRNA 3 | TAATACGACTCACTATAGGG tccactttcctcccatctgc GTTTTAGAGCTAGAAATAG |
| fwd_mouse *Cer1* gRNA 4 | TAATACGACTCACTATAGGG ggatgactccctggaacgcc GTTTTAGAGCTAGAAATAG |
| fwd_mouse *Gsc* gRNA 1 | TAATACGACTCACTATAGGG cgcgtctttgcagcgcggcc GTTTTAGAGCTAGAAATAG |
| fwd_mouse *Gsc* gRNA 2 | TAATACGACTCACTATAGGG gactcgctctacggcgccgg GTTTTAGAGCTAGAAATAG |
| fwd_mouse *Gsc* gRNA 3 | TAATACGACTCACTATAGGG agctgccgaccgcggccggg GTTTTAGAGCTAGAAATAG |
| fwd_mouse *Gsc* gRNA 4 | TAATACGACTCACTATAGGG cacgtgcaggcggcgcccgt GTTTTAGAGCTAGAAATAG |
| fwd_mouse *Six3* gRNA1 | TAATACGACTCACTATAGGG gttgatggcctcgcacgccc GTTTTAGAGCTAGAAATAG |
| fwd_mouse *Six3* gRNA2 | TAATACGACTCACTATAGGG gtacaggtcgcggaagttgc GTTTTAGAGCTAGAAATAG |
| fwd_mouse *Six3* gRNA3 | TAATACGACTCACTATAGGG gtggctcgaggcgcactacc GTTTTAGAGCTAGAAATAG |
| fwd_mouse *Six3* gRNA4 | TAATACGACTCACTATAGGG ggccagcgtctgcgagacgc GTTTTAGAGCTAGAAATAG |
| fwd_mouse *Hesx1* gRNA 1 | TAATACGACTCACTATAGGG attattctaggtcgaagtat GTTTTAGAGCTAGAAATAG |
| fwd_mouse *Hesx1* gRNA 2 | TAATACGACTCACTATAGGG ccctggcattgacatcagag GTTTTAGAGCTAGAAATAG |
| fwd_mouse *Hesx1* gRNA 3 | TAATACGACTCACTATAGGG aactgtgtttgcaccctacc GTTTTAGAGCTAGAAATAG |
| fwd_mouse *Hesx1* gRNA 4 | TAATACGACTCACTATAGGG gaggaagacagaatccaggt GTTTTAGAGCTAGAAATAG |

fwd: forward primer; rev: reverse primer; OF: outer forward primer; OR: outer reverse primer; IF: inner forward primer; IR: inner reverse primer. The gRNA sequence is marked in red

Primer used for genotyping of targeting site.

| Name | Sequence (5'→3') |
|---|---|
| *Pdx1* Genotyping F | ctccacagcagcaagcagggatcagg |
| *Pdx1* Genotyping R | atggagaacaagctgccaacatgagtggac |
| *Pax6* Genotyping OF | cccatgcagatgcaaaagtcc |
| *Pax6* Genotyping OR | ctttgacccctggtctgtgg |
| *Pax6* Genotyping IF | gatgcaaaagtccaggtgct |
| *Pax6* Genotyping IR | cccaggtaccctggagacaa |
| *Lhx1* Genotyping OF | gcccgcttgagactgatttt |
| *Lhx1* Genotyping OR | ctgaagccatctcagaccct |
| *Lhx1* Genotyping IF | ccctctaccagcctggtctctccccc |
| *Lhx1* Genotyping IR | ccggaccacgctttccagaaccaagtac |
| *Otx2* Genotyping OF | ccgcattacactgggaggtc |
| *Otx2* Genotyping OR | actgtagggactcttgcgac |
| *Otx2* Genotyping IF | gtggggatactgggtaacgc |
| *Otx2* Genotyping IR | gggactcttgcgacctacag |
| *DKK1* Genotyping OF | ctgcttccgacacacaaaca |
| *DKK1* Genotyping OR | gagcaactcaagcatcctgg |
| *DKK1* Genotyping IF | gaaatcccatcccggctttg |
| *DKK1* Genotyping IR | actcaagcatcctggtaccc |
| *Cer1* Genotyping OF | gagcctctcttttaggcccg |
| *Cer1* Genotyping OR | atgtgcttacatttggccacg |
| *Cer1* Genotyping IF | agcctctcttttaggcccgt |
| *Cer1* Genotyping IR | ggaactagacaaggcccgag |
| *Gsc* Genotyping OF | cgcgctctctttcggtttg |
| *Gsc* Genotyping OR | cgtcctcacttcattcccgc |
| *Gsc* Genotyping IF | gctctctttcggtttgctcg |
| *Gsc* Genotyping IR | agacgtatgcaaagtccccg |
| *Six3* Genotyping OF | tcagtccatggtattccgct |
| *Six3* Genotyping OR | cgccggttcttaaaccagttg |
| *Six3* Genotyping IF | tcagtccatggtattccgctc |
| *Six3* Genotyping IR | aatgggtcttctgctcgcc |
| *Hesx1* Genotyping OF | tgagttggtaccgaggacga |
| *Hesx1* Genotyping OR | gggagaatggggtgagtctg |
| *Hesx1* Genotyping IF | ggtcccagtgtagaagtggc |
| *Hesx1* Genotyping IR | agtgttaggggaggaacgga |
